# A scalable MNase-seq framework for reproducible nucleosome profiling across pluripotent stem cell and cardiomyocyte models

**DOI:** 10.64898/2026.06.08.731013

**Authors:** Chris Thekkedam, David T. Humphreys, Marina Naval-Sanchez, Amy M. Nicks, Richard P. Harvey, Osvaldo Contreras

## Abstract

Micrococcal nuclease (MNase) digestion is widely used to profile chromatin accessibility and nucleosome footprinting. However, its application is often limited by sensitivity to reaction conditions, high cell input requirements, and the lack of standardized protocols across cell types. Here we developed a robust MNase workflow encompassing buffer composition, DNA purification chemistry, fixation and decrosslinking parameters, cell input scalability, and an in-house yeast spike-in for quantitative normalization. We validated this unified framework across human induced pluripotent stem cells (hiPSCs), hiPSC-derived cardiomyocytes at multiple differentiation stages, primary murine embryonic cardiac cells, and adult mouse cardiomyocytes, and demonstrated comparable digestion efficiencies and kinetics despite marked differences in cellular architecture and chromatin organization. Genome-wide MNase-seq in hiPSCs, combined with the *nucMACC* bioinformatic pipeline, resolved concentration-dependent nucleosomal occupancy and precise nucleosome positioning at pluripotency-related regulatory elements. This modular, end-to-end, and scalable workflow provides a standardized platform for reproducible MNase-based chromatin profiling across diverse *in vitro* and *in vivo* models.

**TEASER:** A unified, rapid, and scalable MNase-seq workflow for reproducible mononucleosomal and subnucleosomal profiling from stem cells to adult cardiomyocytes.

**HIGHLIGHTS:**

- Systematic MNase optimization across buffer, DNA purification, and cell input variables
- Unified workflow validated in hiPSCs, hiPSC-CMs, embryonic, and adult cardiomyocytes
- Fixed-cell protocol enables weeks of storage without loss of DNA quality
- Cost-effective yeast spike-in ensures quantitative normalization for MNase-seq
- Genome-wide analyses confirm robust and precise nucleosome positioning at regulatory elements

## INTRODUCTION

Nucleosome positioning and chromatin dynamics are fundamental determinants of DNA accessibility and the regulation of transcription, DNA replication, and repair [1,2]. Nucleosomes, the basic histone–DNA units of chromatin, are highly dynamic structures whose organization is shaped by DNA sequence, chromatin remodelers, and transcriptional machinery, collectively generating a continuously evolving chromatin landscape [1,3–5]. By controlling which genomic regions are accessible to regulatory proteins, nucleosome organization influences transcription factor binding, RNA polymerase progression, and cell-type-specific gene expression programs. Understanding how chromatin landscapes are established and remodeled is therefore central to fields ranging from developmental biology to disease modeling [2,6,7].

Micrococcal nuclease sequencing (MNase-seq) has become a principal technique for interrogating nucleosome organization genome-wide [5,7–14]. It exploits MNase’s propensity to selectively digest accessible linker DNA while preserving nucleosome-bound DNA [15,16], releasing mononucleosomal fragments whose genomic distribution reveals nucleosome occupancy and positioning [4,15,17–20]. Unlike open-chromatin assays such as DNase-seq and ATAC-seq (Assay for Transposase-Accessible Chromatin using sequencing), MNase-seq enables isolation and sequencing of nucleosome core particles, directly profiling nucleosome-occupied DNA [8,10,12,21]. New computational frameworks such as *nucMACC* leverage differential MNase digestions to quantify single-nucleosome stability and accessibility *in situ* and at single-nucleosome resolution, moving beyond nucleosome occupancy [17,22]. For example, ultra-deep paired-end MNase-seq mapping showed that much of the apparent nucleosome “fuzziness” in earlier low-depth maps is resolved at higher sequencing coverage, revealing far more precisely positioned nucleosomes across cell populations [23]. Collectively, these advances highlight nucleosome dynamics as an additional layer of regulatory information tied to transcriptional potential and activity [4,5,22,24–26].

However, broad adoption of MNase-based chromatin profiling remains constrained by technical variability and a lack of standardization. These limitations stem in part from the enzyme itself: MNase preferentially cleaves adenine/thymine (A/T)-rich DNA—a sequence bias shared across other chromatin accessibility assays [8,15,27]—introducing systematic distortions into nucleosome positioning estimates [19,21,28,29], while its dual endo- and exonuclease activities risk degrading nucleosomal DNA at excessive digestion levels, obscuring occupancy signals at heavily digested loci [12,18]. Beyond enzyme-specific factors, experimental variables in buffer composition, DNA purification, cell input, and fixation conditions can all substantially influence the resolution and reproducibility of resulting MNase-seq profiles. This variability is further compounded by the absence of commercial kits for MNase-based workflows, meaning that most laboratories develop protocols empirically for a single cell type. Moreover, the absence of unified workflows complicates cross-study comparisons and integration of MNase-derived datasets with emerging quantitative omics platforms; a challenge exacerbated by insufficient methodological details in many published MNase-seq studies [17,19,28–30], and that ultimately hinders reproducibility across laboratories.

Another practical challenge is the need for quantitative cross-sample normalization. Because MNase digestion efficiency varies with enzyme concentration, incubation time, and chromatin context, direct comparisons of nucleosomal profiles across samples require an exogenous DNA reference standard. While spike-in DNA can fulfil this role [31,32], such reagents impose substantial per-sample costs and are subject to batch variability, limiting their utility in large experiments.

In the cardiovascular context, these challenges are particularly acute. Nucleosome dynamics during cardiomyocyte differentiation and maturation are underexplored. Human induced pluripotent stem cell-derived cardiomyocytes (hiPSC-CMs) have emerged as a powerful platform for modeling cardiovascular development and disease [33–35]. However, MNase-based chromatin assays in these cells are complicated by their mechanical fragility, heterogeneity, and need for cardiomyocyte enrichment prior to analysis. These barriers are exacerbated in primary cardiac tissue. Adult mouse cardiomyocytes yield limited cell numbers and are large, often binucleated cells with a rigid sarcomeric architecture, making chromatin preparation exceptionally challenging.

Here, we present a systematically optimized *in vivo* MNase chromatin digestion workflow that can be completed within half a day, addressing these challenges. By optimizing key experimental parameters—including buffer composition, DNA purification, enzymatic conditions, and cell input—we establish a robust, reproducible framework for nucleosomal profiling, in line with the *PRO□MaP* best-practice guidelines for transparent methodology [36]. To enable accurate cross-sample comparisons without reliance on commercial reagents, we implement a cost-effective in-house spike-in calibration strategy using yeast. Genome-wide MNase-seq in hiPSCs, coupled with *nucMACC* bioinformatic analysis, shows that our optimized conditions resolve concentration-dependent shifts in nucleosome accessibility and occupancy with high precision at regulatory elements. Orthogonal validation against independent Encyclopedia of DNA Elements (ENCODE) ATAC-seq, DNase-seq, and histone modification ChIP-seq datasets confirms that *nucMACC*-classified nucleosome accessibility states correspond to established regulatory chromatin landscapes, providing external cross-platform support for the biological fidelity of our workflow. Finally, we further validate this MNase-based framework using a spectrum of cardiac models of increasing complexity, including hiPSC-derived cardiomyocytes at multiple differentiation stages, primary embryonic murine cardiac cells, and adult mouse cardiomyocytes.

## RESULTS

### Systematic optimization of an MNase-based chromatin digestion workflow

Although MNase digestion remains a cornerstone of chromatin biology and nucleosomal footprinting, its utility is often limited by sensitivity to buffer composition, enzyme calibration, lack of standardized reagents, protocol length, and variable DNA recovery efficiency. To address these challenges, we optimized a custom MNase chromatin digestion workflow by systematically evaluating buffer composition, digestion chemistry, enzymatic treatments, DNA purification, and cell input scalability. An overview of the optimized workflow is shown in **Fig. 1A**. Briefly, cells are lysed to release intact chromatin, followed by Ca²-dependent MNase chromatin digestion to generate nucleosomal fragments. Fragmented DNA is then purified and resolved by electrophoresis to assess nucleosomal ladder quality.

**Figure 1.**
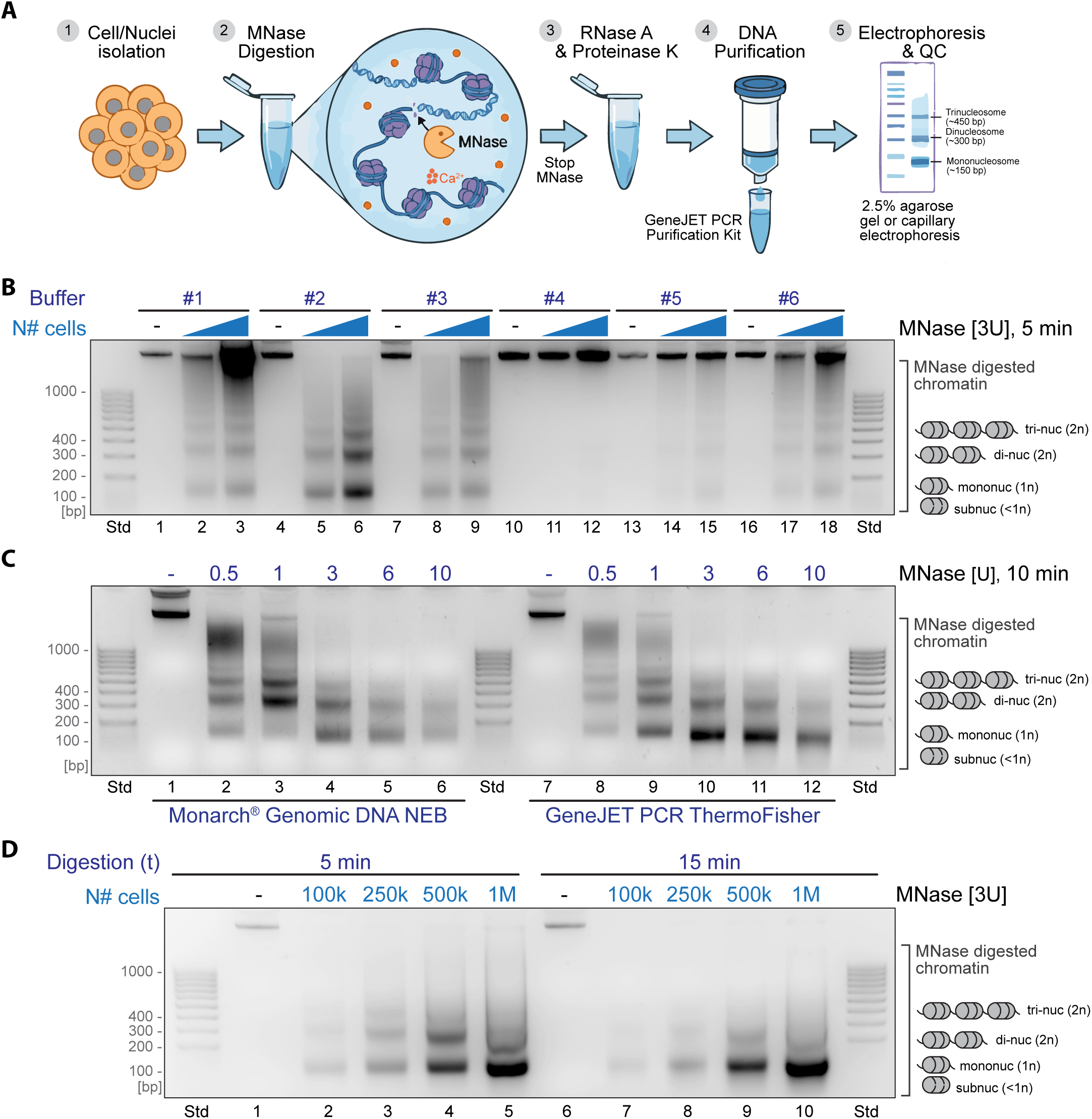
Standardization of buffers, DNA purification kits, and cell numbers for MNase-mediated digestion of chromatin. (A) Schematic overview of the MNase-seq sample preparation and validation workflow. The protocol is divided into five chronological steps: (1) Cell/Nuclei isolation: Starting material consisting of cells or intact nuclei is prepared. (2) MNase Digestion: Chromatin is incubated with Micrococcal Nuclease (MNase) and calcium ions (Ca^2+^). MNase preferentially cleaves the accessible linker DNA between histone octamers, generating nucleosome-protected DNA fragments. (3) RNase A & Proteinase K Treatment: The enzymatic reaction is terminated. RNase A is added to degrade contaminating RNA, followed by Proteinase K to digest histones and other chromatin-associated proteins, releasing the DNA. (4) DNA Purification: The fragmented DNA is isolated and cleaned using a spin-column based method (e.g., GeneJET PCR Purification Kit). (5) Electrophoresis & Quality Control (QC): Purified DNA is visualized via 2.5% agarose gel or capillary electrophoresis. The schematic illustrates a characteristic nucleosomal ladder, with distinct bands corresponding to mononucleosomes (approx. 147 bp), dinucleosomes, and trinucleosomes, confirming successful chromatin digestion and high-quality DNA prior to library preparation. (B) 2.5% agarose gel electrophoresis was used to analyze MNase digestion products, comparing two distinct buffers (Buffer 1: B#1 vs. Buffer 2: B#2) and their efficiency when loading the same volume of digested chromatin. The GeneJET PCR Kit was used to isolate and purify digested DNA, with 500,000 hiPSCs used in all lanes. Buffer 2 demonstrated superior discrimination of all nucleosomal fractions compared to Buffer 1. (C) In an MNase titration experiment (10 minutes incubation), 2.5% agarose gel electrophoresis was used to analyze MNase digestion products, comparing two DNA isolation and purification kits. 500,000 hiPSCs were used. The GeneJET PCR Kit from ThermoFisher demonstrated superior discrimination of mono-nucleosomal fractions compared to the Monarch® Genomic DNA Kit from NEB. (D) MNase digestion products were analyzed using different cell numbers with the GeneJET PCR Kit from ThermoFisher. Undigested genomic DNA (lanes 1 and 6) was purified from 100,000 cells (100k).

Buffer composition is a primary determinant of MNase digestion efficiency, yet most published protocols adopt a single formulation without a comparative analysis. To define optimal conditions for nucleosomal profiling in hiPSCs, we systematically adapted and evaluated six published buffer formulations (see **Methods**). Buffer 2 (B#2) produced superior nucleosomal ladder resolution relative to all other conditions, yielding higher DNA recovery and clearer separation of mono-, di-, and tri-nucleosomal fragments (**Fig. 1B**). Buffers B#1 and B#3 performed comparably well, whereas B#5 and B#6 underperformed, and B#4 exhibited the lowest DNA digestion and recovery (**Fig. 1B**).

We next evaluated the impact of DNA purification chemistry on fragment recovery. Using an MNase titration (0.5–10 U) across two commercial DNA purification systems under identical conditions, we observed marked differences in performance. While the Monarch Genomic DNA Kit (NEB) provided adequate recovery, the GeneJET PCR Kit (ThermoFisher) yielded substantially improved resolution and DNA quantity of mono- and subnucleosomal fragments, particularly at higher enzyme concentrations (1–10 U) (**Fig. 1C**). Surprisingly, the GeneJET system also enabled clearer discrimination of the mononucleosomal-to-dinucleosomal and mononucleosomal-to-trinucleosomal ratios, a critical metric for assessing digestion quality, with optimal digestion typically corresponding to ∼70–80% mononucleosomes and 20–30% higher-order nucleosomal fragments [8,37,38].

A major limitation of mammalian MNase-based assays is the requirement for high cell number input. To assess scalability, we applied the optimized workflow across inputs ranging from 100,000 to 1 × 10□ hiPSCs. Nucleosomal profiles remained highly consistent across this range, with stable digestion kinetics observed at both 5- and 15-minute incubations (**Fig. 1D**). Notably, high-resolution fragment discrimination was retained even at low cell inputs without additional protocol modifications. Collectively, these findings demonstrate that our workflow is both scalable and reproducible in hiPSCs.

### Kinetic characterization and dose-dependent control of chromatin fragmentation

To achieve the precise fragment size distribution required for high-resolution MNase profiling, we next sought to characterize the digestion kinetics of MNase in hiPSCs. Because the transition from bulky chromatin to discrete nucleosomal units is a dynamic process [17,19,39], we benchmarked the workflow across a wide range of enzyme concentrations and incubation times to establish conditions that allow users to tailor fragmentation to their experimental needs.

We first performed a comprehensive dose-response titration, varying MNase concentrations from 0.5 U to 50 U over a fixed 10-minute interval using 500,000 cells (**Supplementary Fig. 1A**). Our results demonstrated a highly predictable transition in chromatin architecture: at lower concentrations (0.5–1 U), genomic DNA remained largely associated with higher-order oligonucleosomes, whereas intermediate doses (3–6 U) generated the characteristic nucleosome ladder, cleanly resolving mono- (1n), di- (2n), and tri-nucleosomal (3n) fragments. At the highest concentrations (10–50 U), digestion yielded predominantly mononucleosomal and subnucleosomal fragments—a prerequisite for high-sensitivity MNase-seq applications (sometimes termed “overdigestion”) [17,22,39] and that ultimately undermines reproducibility. This dose–dependence underscored the robustness of the system, allowing researchers to calibrate chromatin digestion based on their samples and cell models to reproducibly generate a high yield of mononucleosomal fragments.

Recognizing that MNase enzyme concentration and time are interdependent variables, we conducted a time-course analysis comparing two enzyme concentrations: 1 Unit and 3 Units (**Supplementary Fig. 1B**). With 1 Unit of MNase, the transition to mononucleosomal (1n) fragments was gradual, maintaining a balanced distribution of 2n and 3n species even after 30 minutes. Conversely, the 3-Unit condition accelerated this trajectory, achieving a predominantly mononucleosomal state within 5 minutes. These kinetic profiles demonstrated that our optimized protocol offers precise control over chromatin fragmentation and enables reliable fragmentation.

### Optimization of PFA fixation and crosslink reversal for fixed-cell MNase digestion

Although digestion of native chromatin is highly efficient, many epigenomic workflows require formaldehyde (PFA) fixation to preserve chromatin structure and protein–DNA interactions. This is particularly relevant for low-input applications such as single-cell MNase-seq (scMNase-seq) [40,41] and cross-linking chromatin immunoprecipitation followed by sequencing (X-ChIP-seq). Fixation confers two key advantages for chromatin studies: it stabilizes the chromatin landscape, preventing artifactual nucleosome repositioning during sample processing, and it covalently crosslinks histones to DNA, ensuring that MNase digests only accessible linker DNA while nucleosome-bound regions remain protected. Additionally, fixation enables batch processing of samples collected across multiple time points or experimental conditions—a flexibility that is often impractical with native chromatin. However, fixation introduces an inherent trade-off: the covalent crosslinks that preserve chromatin architecture also impede enzymatic accessibility, necessitating controlled crosslink reversal before MNase can effectively digest the substrate. To address this, we systematically optimized PFA fixation and thermal decrosslinking parameters, evaluating storage stability and the efficiency of crosslink reversal (**Supplementary Fig. 2**), validated across different cell types presented in subsequent sections.

A primary concern in fixed-cell assays is the potential for sample degradation or progressive over-crosslinking during prolonged storage. We first benchmarked the performance of our workflow using 1% PFA-fixed hiPSCs stored for two to four weeks at 4°C. The nucleosomal profiles obtained from stored, fixed cells were indistinguishable from those of freshly dissociated cells, maintaining clear fragment resolution across both 250k and 500k cell inputs (**Supplementary Fig. 2A**).

The accessibility of MNase to fixed chromatin is critically dependent on reversal of formaldehyde-induced crosslinks [42], yet standardized decrosslinking protocols for MNase-based assays are lacking. We evaluated the known effect of sample heating at 55°C to facilitate crosslink reversal. Titration of heating durations (10 minutes to 1 hour) revealed that a 10-to-30-minute incubation at 55°C is optimal for restoring fragment resolution, effectively reversing 1% PFA-induced crosslinks while preserving DNA integrity (**Supplementary Fig. 2B–C**). Notably, 55°C heating for one hour or longer (up to 18 h) resulted in a substantial reduction in digested DNA yield (**Supplementary Fig. 2C**), confirming our previous results. This is in contrast to previous studies that employed prolonged overnight heating as a decrosslinking step [25]. Furthermore, we evaluated the synergy between thermal decrosslinking and chemical quenching (glycine) [42] across varying PFA concentrations (0.1% to 2%) (**Supplementary Fig. 2D**). We observed that while 125 mM glycine alone is insufficient to fully rescue digestion at any given PFA concentration, short heating at 55°C effectively restores MNase accessibility, ensuring clean fragment separation of oligonucleosomes (**Supplementary Fig. 2D**). Together, these parameters define a standardized set of conditions for reproducible chromatin fragmentation from fixed cells.

### Development and validation of a cost-effective, in-house yeast spike-in for quantitative normalization

Accurate quantitative comparison across MNase-seq libraries is frequently hampered by the cost and limited flexibility of commercial spike-in reagents. To overcome these barriers, we developed a streamlined, in-house pipeline for generating a robust *Saccharomyces cerevisiae* spike-in DNA from log-phase cultures, providing a robust internal control for downstream sequencing applications (**Supplementary Fig. 3A**) [31,32].

We first optimized the enzymatic recovery of yeast mononucleosomes by subjecting spheroplasted cells (OD ≈ 0.9) to high-dose MNase digestion (60 U for 10 minutes). To determine the optimal DNA input for library preparation, we benchmarked two cell quantities (see **Methods**). Gel electrophoresis of the digested DNA revealed that the 1× input yielded a substantially more abundant and discrete mononucleosomal fraction compared to the 0.5× cell dilution, making it the preferred condition for spike-in calibration (**Supplementary Fig. 3B**). This in-house pipeline enables the generation of large-scale, consistent spike-in batches, circumventing the lot-to-lot variability often associated with commercial reagents.

A critical requirement for any spike-in reagent is efficient incorporation into library preparation and sequencing workflows [31,32]. To validate the performance of our yeast-derived fragments, we integrated them at 5% of total isolated DNA into hiPSC samples subjected to a broad MNase titration gradient (0.5–10 U). Next-generation paired-end sequencing analysis revealed that the spike-in mononucleosomes exhibited invariant fragment density and consistent footprinting patterns relative to the transcriptional start site (TSS) across all digestion conditions (**Supplementary Fig. 3C**). This consistency confirms that the yeast spike-in DNA maintains its structural integrity and recovery profile regardless of the host sample’s digestion state, validating it as a reliable standard for quantitative cross-sample normalization.

### Concentration-dependent changes in sub- and mononucleosome occupancy revealed by next-generation sequencing

To extend our bench-top optimization to a genome-wide scale, we integrated the workflow into the recently published *nucMACC* bioinformatic pipeline [22] (**Fig. 2A**). *nucMACC* provides a genome-wide strategy for quantifying nucleosome accessibility and stability at single-nucleosome resolution, extending MNase-seq beyond conventional occupancy mapping. Capillary electrophoresis of in-house-prepared NGS libraries confirmed that the optimized digestion conditions produced high-quality libraries compatible with both NextSeq 50-bp paired-end and NovaSeq-X 150-bp paired-end sequencing platforms (**Fig. 2B–C**).

**Figure 2.**
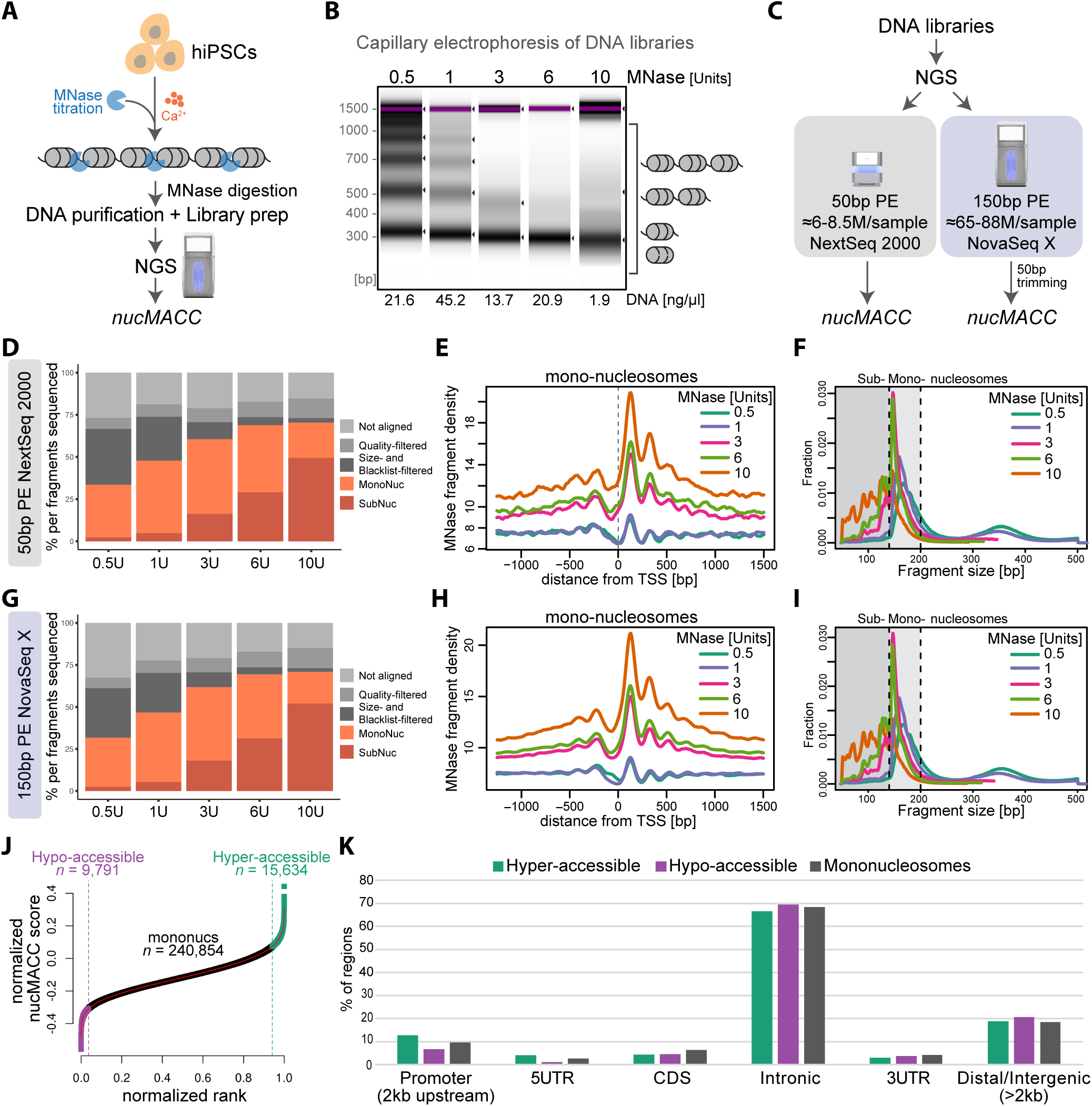
MNase concentration-dependent changes in sub-nucleosome and mono-nucleosome accessibility. (A) Experimental outline of our hiPSC MNase-seq tritation experiment and nucMACC analysis. (B) Capillary electrophoresis of DNA libraries used for NGS sequencing of digestion products from our MNase titration experiment using 250,000 hiPSCs. (C) Experimental outline describing the two NGS strategies and later nucMACC analysis. (D) Proportion of DNA fragments under increasing MNase concentrations, showing a reduction in mono-nucleosomal fractions and an increase in sub-nucleosomal fractions. Data was obtained from 50bp paired-end NGS. (E) Average fragment frequencies at the transcriptional start site (TSS) in base pairs (bp). (F) Fragment size distribution of MNase titrations. Fragments are selected based on size and grouped into sub-nucleosomes (<140 bp, dark grey section) or mono-nucleosomes (140 to 200 bp, light grey section). (G-I) Same description as per (D-F) with data obtained from 150bp PE NGS. (J) Selection of hyper-accessible (green) and hypo-accessible (purple) nucleosomes based on the nucMACC scores. Nucleosomes were ranked by the nucMAcc score. x and y axes were normalized to a range of one. Cutoffs were deduced from the slope of the LoeSS curve fit. Nucleosomes before the point, where the slope goes the first time below 1 (purple dashed line), were classified as hypo-accessible. Nucleosomes after the point where the slope of the curve exceeds one again (green dashed line), were classified as hyper-accessible nucleosomes. (K) Distribution of the hypo-/hyper-accessible nucleosomes and mononucleosomes peaks relative to gene features in the human genome. Peaks were classified according to their co-location with gene features as, transcription start site (TSS), 5’ untranslated region (5’UTR), 3’UTR, promoter, coding sequence (CDS), intronic and intergenic.

Sequencing analysis of hiPSCs using *nucMACC* revealed a clear, dose-dependent transition in fragment populations. Increasing MNase concentration progressively enriched for subnucleosomal fragments (<140 bp) while depleting the mononucleosomal fraction (**Fig. 2D, G**). When mapped around transcription start sites (TSSs), these digestion-dependent changes resolved the nucleosome-depleted region (NDR) and the positioning of the +1 and −1 nucleosomes with high precision (**Fig. 2E, H**). This pattern is consistent with canonical promoter architecture, in which an accessible NDR is flanked by well-positioned nucleosomes that define transcriptional regulatory boundaries [1,5,25,43].

Notably, higher MNase conditions, particularly 3–10 U, maximized accessibility contrast at these regulatory interfaces (**Fig. 2E, H**), indicating that tuning the MNase-to-cell ratio enables fine control over the resolution of chromatin accessibility maps. At the same time, increasing MNase concentration consistently shifted the fragment distribution toward smaller subnucleosomal species, demonstrating that higher MNase units enhance accessibility detection but also increase the generation of subnucleosomal fragments (**Fig. 2D–I**). In addition, genome-wide correlation of mononucleosomal read counts confirmed that higher MNase concentrations (3–10 U) produced profiles highly concordant with published iPSC MNase-seq datasets (*Pearson’s r* = 0.86–0.93) [25], whereas lower concentrations (0.5–1 U) diverged substantially (*r* = 0.20–0.39), further supporting 3–6 U as the optimal digestion range and validating our study (**Supplementary Fig. 4A**).

To move beyond bulk occupancy and resolve accessibility at the level of individual nucleosomes, we used the *nucMACC* score—calculated as the slope of a linear regression across MNase titration conditions and normalized for local GC content to correct for MNase sequence bias (**Supplementary Fig. 4B**)—to stratify mononucleosomes into discrete accessibility classes. The full set of mononucleosomes (*n* = 266,279 nucleosomes) was ranked by normalized *nucMACC* score, the rank–score relationship was scaled to a unit range, and class boundaries were derived from the geometry of a LOESS fit rather than from arbitrary thresholds. Nucleosomes preceding the point at which the local slope first fell below one were classified as *hypo-accessible* (*n* = 9,791), and those beyond the point at which the slope again exceeded one as *hyper-accessible* (*n* = 15,634) (**Fig. 2J**). This data-driven cutoff isolates the two tails of the accessibility spectrum that depart most strongly from the bulk population.

We next asked whether these classes occupy distinct positions within gene architecture. Annotating hypo-accessible, hyper-accessible, and bulk mononucleosome peaks by their co-location with genomic features showed that all three populations were most frequently intronic (∼64–68%) and, secondarily, distal/intergenic (∼18–20%), consistent with the large genomic coverage of these compartments (**Fig. 2K**). The classes nonetheless partitioned distinctly: hyper-accessible nucleosomes were markedly over-represented at promoters (2 kb upstream) and 5′UTR relative to both hypo-accessible and bulk nucleosomes, whereas hypo-accessible nucleosomes showed relatively greater intronic and distal intergenic representation compared to hyper-accessible nucleosomes; CDS and 3′UTR compartments were sparsely occupied across all classes (**Fig. 2K**). This preferential promoter positioning of the most accessible nucleosomes indicates that *nucMACC* scores capture a genuine regulatory organization of the genome, providing the foundation for the orthogonal validation presented below.

### nucMACC-defined nucleosome accessibility states correspond to distinct regulatory chromatin landscapes in pluripotent stem cells

We next asked whether *nucMACC*-defined mononucleosome accessibility corresponds to regulatory chromatin states detected by orthogonal accessibility assays. To address this, we profiled independent genome-wide ATAC-seq and DNase-seq signal around *nucMACC*-classified mononucleosome centers, including hyper-accessible, hypo-accessible, and all mononucleosome positions (**Fig. 3A**). The center of each profile corresponds to the midpoint of the *nucMACC* mononucleosome interval, rather than a TSS or ATAC/DNase peak summit. Thus, hyper- and hypo-accessible classes refer to nucleosome positions with high or low MNase-derived accessibility scores, not to accessibility classes defined by ATAC-seq or DNase-seq.

**Figure 3.**
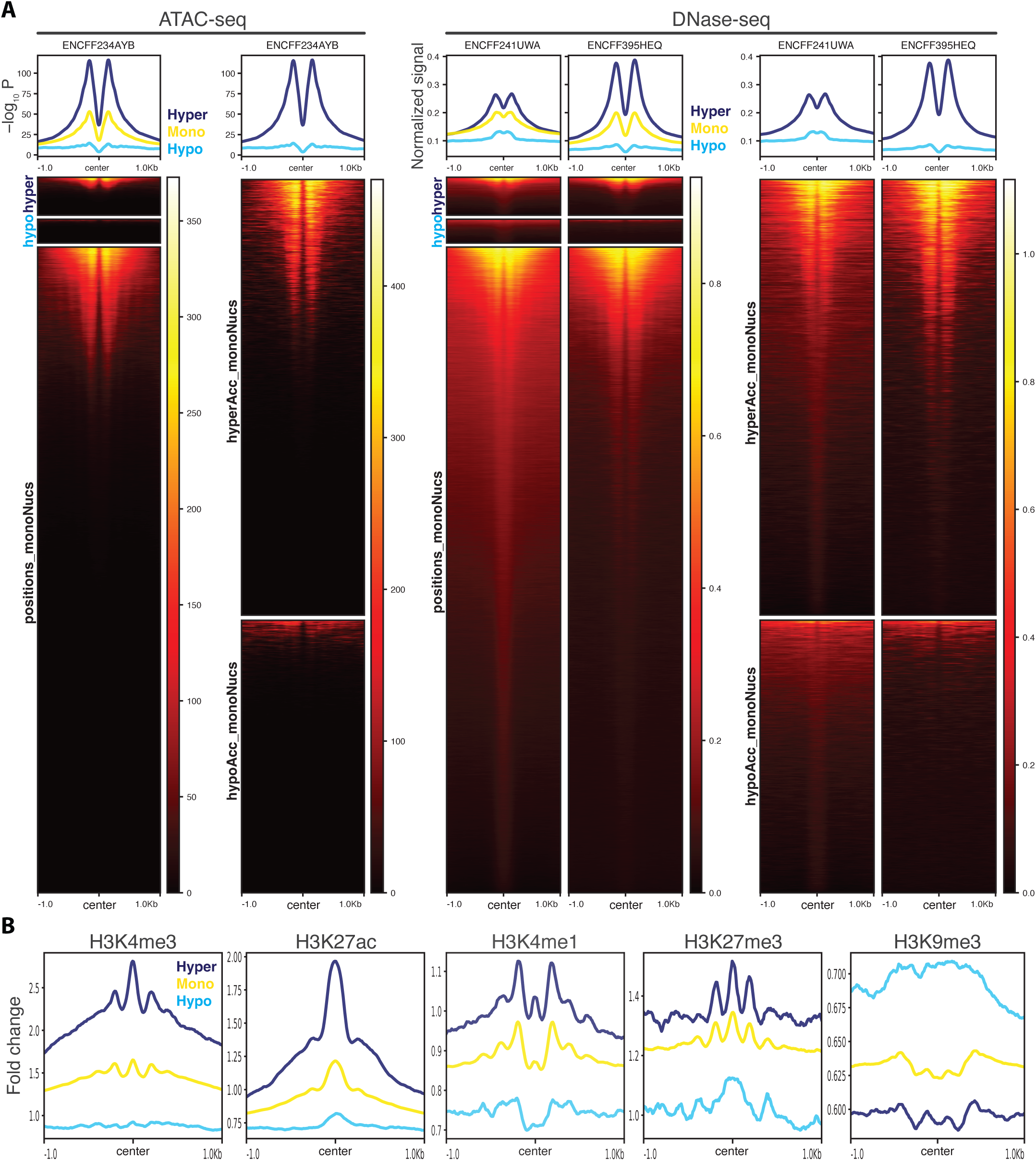
nucMACC-derived nucleosome accessibility states are concordant with independent chromatin accessibility and histone modification profiles in pluripotent stem cells. (A) Average profiles (top) and heatmaps (bottom) of ENCODE hiPSC ATAC-seq and DNase-seq centered on MNase-seq nucleosome dyads, stratified by nucMACC class: hyper-accessible (navy), bulk mononucleosomes (yellow), and hypo-accessible (cyan). The y-axis shows mean signal intensity across regions (ATAC-seq, ENCODE −log₁₀ P; DNase-seq, read-depth-normalized signal). Hyper-accessible nucleosomes show stronger dyad signal and well-phased flanking periodicity, indicating regular nucleosome arrays at accessible regions, whereas hypo-accessible nucleosomes show reduced signal and weaker periodicity, consistent with compacted chromatin. Concordance across both assays supports the nucMACC classification. (B) Average ChIP-seq enrichment for H3K4me3, H3K27ac, H3K4me1, H3K27me3, and H3K9me3 centered on the same three classes, shown as fold-change over control (input). Active and/or poised marks (H3K4me3, H3K27ac, H3K4me1, H3K27me3) are enriched at hyper-accessible nucleosomes, consistent with pluripotent regulatory-element identity, whereas a repressive mark (e.g., H3K9me3) is enriched at hypo-accessible nucleosomes.

Hyper-accessible mononucleosomes showed the strongest enrichment of ATAC-seq and DNase-seq signal around their genomic positions, whereas hypo-accessible mononucleosomes showed markedly reduced signal (**Fig. 3A**). The full mononucleosome set displayed an intermediate profile, consistent with a mixed population of accessibility states. This graded relationship was concordant across independent ATAC-seq and DNase-seq datasets, establishing that *nucMACC* accessibility scores reflect genuine differences in the underlying open chromatin landscape [17,22]. To determine whether this stratification extends to the histone modification landscape, we next examined ENCODE [44] ChIP-seq data for the enrichment of five H3 modifications spanning the active-to-repressive chromatin spectrum in hiPSCs.

We profiled H3K4me3, H3K27ac, H3K4me1, H3K27me3, and H3K9me3 centered on the same three nucleosome classes (**Fig. 3B**). The resulting enrichment profiles revealed a striking correspondence between MNase-derived accessibility scores and the histone modification landscape. Marks of active regulatory chromatin partitioned strongly with hyper-accessible mononucleosomes: H3K4me3 and H3K27ac signal peaked sharply at these positions, consistent with their residence at active promoters and enhancer-associated elements [45,46], while concurrent H3K4me1 enrichment further supports their location within enhancer-proximal chromatin [47,48] (**Fig. 3B**). In contrast, H3K9me3, a mark of constitutive heterochromatin [49], showed the inverse pattern, with strongest enrichment at hypo-accessible nucleosomes and minimal signal at hyper-accessible positions, placing hypo-accessible nucleosomes within compacted, transcriptionally silent domains. Notably, H3K27me3 did not follow the canonical repressive distribution: rather than partitioning with hypo-accessible nucleosomes, it was enriched at hyper-accessible positions, co-occurring with H3K4me3 and H3K27ac (**Fig. 3B**). This pattern is consistent with the high prevalence of bivalent and poised promoters in pluripotent cells, where H3K4me3 and H3K27me3 are co-deposited at developmentally regulated loci that remain accessible yet transcriptionally restrained until lineage commitment [50–53]. Bulk mononucleosomes displayed intermediate enrichment across all modifications, reflecting their compositional heterogeneity as an unclassified population spanning both regulatory and structural chromatin (**Fig. 3B**). Together, the coherent stratification of nucleosome classes across both accessibility and histone modification dimensions indicates that *nucMACC* does not merely rank nucleosome signal intensity, but resolves biologically meaningful chromatin states, including the bivalent regulatory architecture characteristic of the pluripotent epigenome.

### Adaptation of the MNase chromatin digestion workflow to hiPSC-CMs

Having established and validated our MNase chromatin digestion workflow using hiPSCs, we next sought to evaluate its applicability in a more complex cellular model. Human iPSC-derived cardiomyocytes represent a powerful platform for modeling cardiac development and cardiovascular disease, yet their use in chromatin profiling studies has been limited by significant technical barriers. Unlike hiPSCs, cardiomyocytes present unique challenges: they are mechanically fragile, difficult to dissociate into viable single-cell suspensions, and are typically obtained as heterogeneous cultures containing non-cardiomyocyte populations. Adapting our workflow for hiPSC-CMs therefore required systematic optimization of cell dissociation and cardiomyocyte enrichment, followed by quality control prior to MNase digestion (**Fig. 4, Supplementary Fig. 5**).

**Figure 4.**
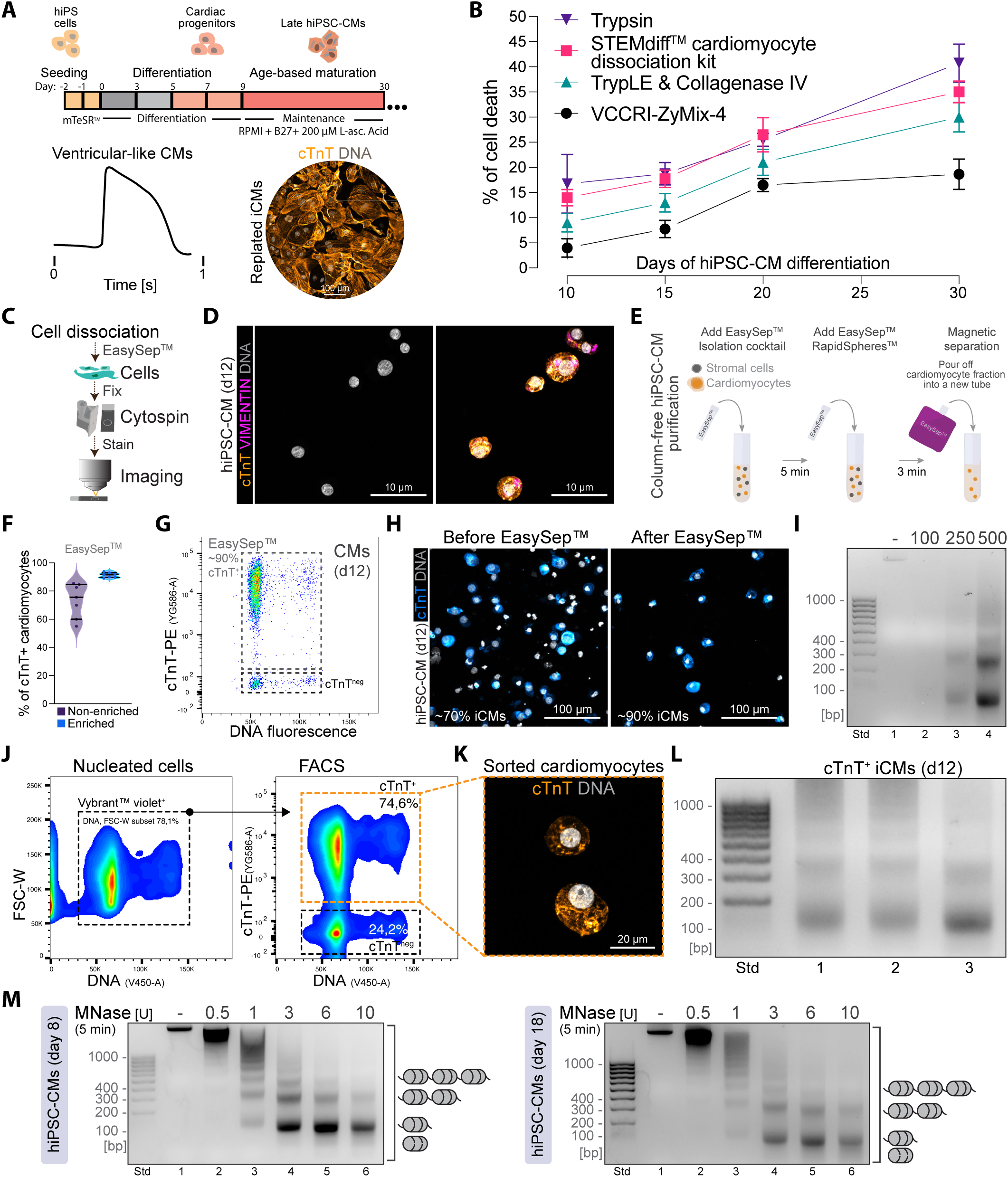
Quality assessment workflow of hiPSC-derived cardiomyocytes for MNase-mediated chromatin digestion. (A) Schematic of the directed differentiation protocol for generating ventricular cardiomyocytes from hiPSCs. Bottom left: representative FluoVolt™ action potential trace (time scale = 1 s). Right: confocal image of replated iCMs at day 22. (B) Cell viability following single-cell dissociation across differentiation stages, comparing four dissociation solutions (n = 4; mean ± SD). VCCRI-ZyMix-4 was selected for downstream applications. (C) Schematic of the post-dissociation quality control workflow: EasySep™ separation, cytospin, staining, and confocal microscopy. (D) Confocal images of day 12 iCMs post-dissociation and cytospin, stained for cTnT, VIMENTIN, and DNA, confirming preserved membrane and nuclear integrity (n = 3). (E) Schematic of column-free EasySep™ cardiomyocyte isolation. (F) Proportion of cTnT+ cardiomyocytes in non-enriched versus EasySep™-enriched fractions across differentiation time points (d8, d12, d18, d28; n = 7). (G) Flow cytometry of the EasySep™-enriched fraction, showing ∼90% cTnT+ cells (n = 2). (H) Confocal images of cytospun day 12 iCMs before and after EasySep™ enrichment, stained for cTnT and DNA (n = 2). (I) MNase digestion (3 U, 5 min) of EasySep™-enriched day 12 iCMs across a range of cell inputs. Lane 1: undigested gDNA from 100,000 cells. (J) Flow cytometry of 1% PFA-fixed day 12 hiPSC-CMs (25,000 cells recorded; n = 3). (K) FACS gating strategy and confocal image of sorted, fixed day 12 iCMs stained for cTnT and DNA (n = 3). (L) MNase digestion (3 U, 2.5 min) of FACS-enriched, fixed day 12 iCMs from three independent sorts (250,000 cells each). (M) MNase titration (0.5–10 U, 5 min) of 250,000 EasySep™-enriched hiPSC-CMs at day 8 and day 18 of differentiation. Lane 1: undigested gDNA from 500,000 cells. (M) MNase titration (0.5, 1, 3, 6 and 10 MNase Units [U]) experiment using 500,000 hiPSC-CMs at day 8 and 18 of directed differentiation. Undigested genomic DNA (lanes 1) was purified from 500,000 hiPSC-CMs.

We first used an in-house WNT directed differentiation protocol to generate ventricular cardiomyocytes from hiPSCs [54], with functional and structural maturation confirmed by *FluoVolt*™ membrane potential recordings and confocal imaging of replated cultures at day 22 post-differentiation (**Fig. 4A**). A critical bottleneck for chromatin studies in cardiomyocyte models is recovering viable, intact single cells. To identify optimal dissociation conditions, we compared four dissociation solutions across multiple stages of cardiomyocyte differentiation, quantifying viability by the proportion of Calcein-AM-positive/propidium iodide-negative cells (*n* = 4). Among the conditions tested, VCCRI-ZyMix-4 consistently yielded the highest cell viability and was therefore selected for all downstream applications (**Fig. 4B**). Confocal microscopy of cytospun cells following dissociation confirmed that VCCRI-ZyMix-4 preserved both cell membrane and nuclear integrity, as assessed by cardiac troponin T (cTnT/TNNT2), VIMENTIN, and DNA staining (**Fig. 4C–D**). Of note, at this stage cardiomyocytes co-express TNNT2/VIMENTIN [55].

Given the heterogeneous composition of hiPSC-CM cultures (i.e., containing stromal cells), we next implemented an EasySep™ magnetic separation step to enrich for cTnT-expressing cardiomyocytes (**Fig. 4E**). Flow cytometry analysis of day 12 directed differentiations (*n* = 7) demonstrated that the enriched fraction contained ≥90% cTnT-positive cells, compared to approximately 75% in non-enriched cultures (**Fig. 4F–G**). This enrichment was corroborated by confocal microscopy of cytospun preparations (**Fig. 4H**). Similarly, we observed high enrichment results when using day 18 hiPSC-CMs (**Supplementary Fig. 5A–C**).

With cell dissociation and cardiomyocyte enrichment conditions established, we assessed whether our optimized MNase chromatin digestion protocol could be directly applied to freshly prepared hiPSC-CMs. Agarose gel electrophoresis of MNase-digested chromatin from EasySep™-enriched day 12 cardiomyocytes (3 U, 5 minutes, 37°C) revealed well-resolved nucleosomal ladders across a range of cell inputs, confirming that the protocol performs robustly in this cell type without further modification (**Fig. 4I**).

We next extended our cell fixation and reverse crosslinking workflow to hiPSC-CMs. Flow cytometry of 1% PFA-fixed day 12 hiPSC-CM cultures was performed followed by FACS-based purification of cardiomyocytes (cTnT+), with the cellular identity of sorted cardiomyocytes confirmed by confocal microscopy. This confirmed efficient cardiomyocyte enrichment and, critically, preservation of cellular and nuclear morphology through fixation and sorting (**Fig. 4J–K**). FACS-enriched, fixed cardiomyocytes (250,000 cells) subjected to decrosslinking and MNase digestion conditions (3 U, 2.5 minutes, 37°C) yielded reproducible nucleosomal profiles across three independent sorting experiments (**Fig. 4L**).

Finally, to determine whether chromatin accessibility changes during cardiomyocyte maturation affect MNase digestion, we performed enzyme titrations (0.5–10 U, 5 minutes) on EasySep™-enriched hiPSC-CMs at day 8 and day 18 of directed differentiation. Both time points produced well-resolved nucleosomal ladders with comparable dose-dependent fragmentation kinetics, indicating that the protocol accommodates the chromatin remodeling associated with cardiomyocyte maturation without requiring stage-specific adjustments (**Fig. 4M and Supplementary Fig. 5A–C**). These results demonstrate that our complete workflow—encompassing hiPSC-CM dissociation, cardiomyocyte enrichment via EasySep™ magnetic or FACS-based separation, fixation, decrosslinking, and MNase digestion conditions—translates effectively from hiPSCs to a substantially more complex differentiated cell type and remains robust across distinct stages of cardiac differentiation.

### In vivo MNase-mediated nucleosomal footprinting of embryonic cardiac cells

Having validated our workflow in hiPSC-CMs, we next asked whether it could be applied to primary cardiac tissue, where cell heterogeneity, extracellular matrix complexity, and limited cell yield present additional challenges. To test this, we isolated hearts from E14.5 mouse embryos and subjected them to enzymatic tissue dissociation [54], cell fixation, and storage at 4°C prior to MNase digestion (**Fig. 5A**). To assess single-cell quality following cardiac tissue dissociation, we performed robust quality control comprising cytospin, cell staining, confocal imaging, and flow cytometry (**Fig. 5A**).

**Figure 5.**
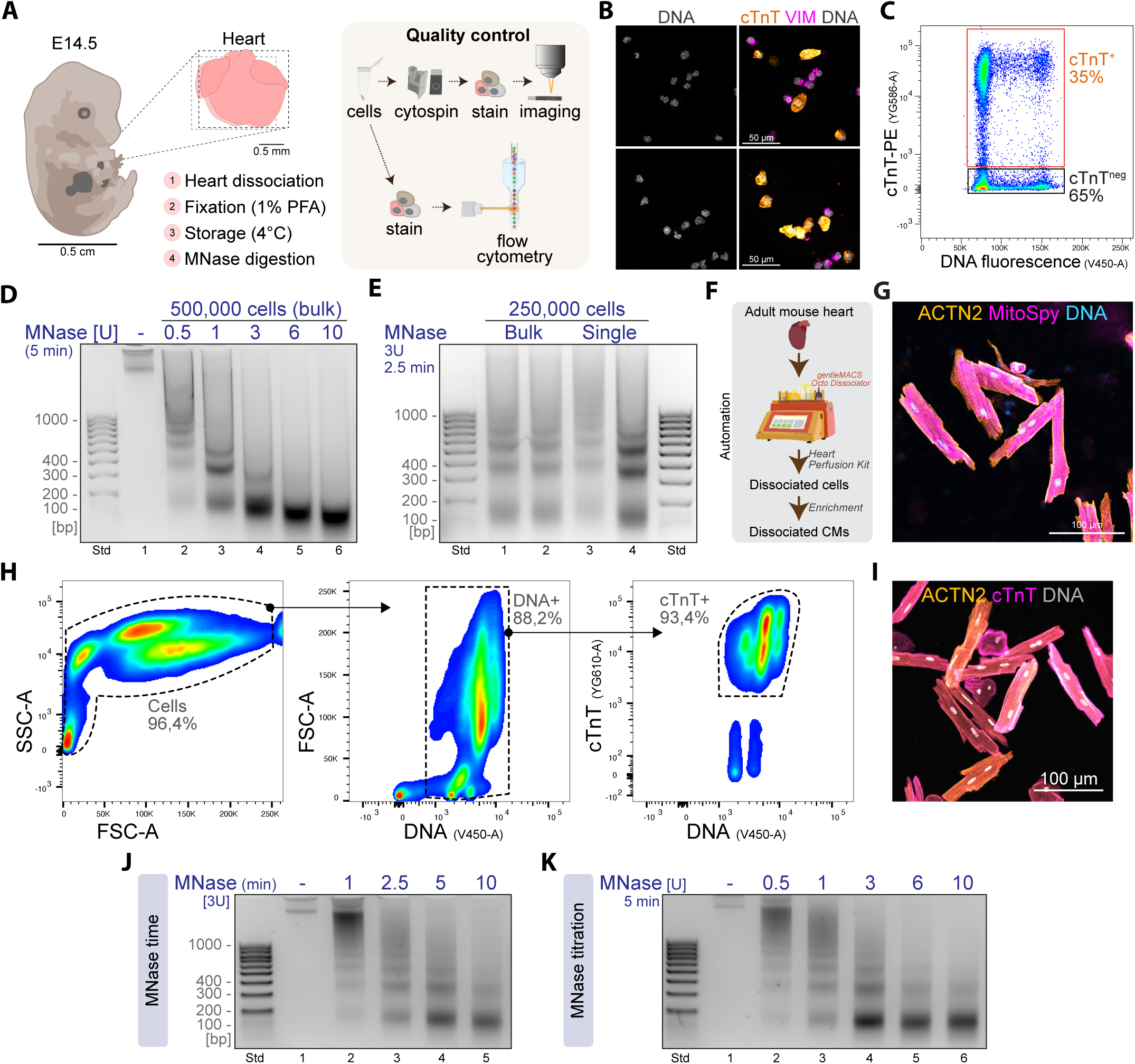
in vivo MNase-mediated nucleosomal footprinting of embryonic cardiac cells and adult cardiomyocytes. (A) Workflow schematic illustrating single-cell dissociation, 1% PFA fixation, 4°C storage, and MNase-mediated chromatin digestion of E14.5 mouse embryonic hearts. Scale bars: 0.5 cm (embryo), 0.5 mm (heart). Quality control pipeline for dissociated E14.5 cardiac cell suspensions, comprising cytospin preparation, immunofluorescence staining, confocal imaging, and flow cytometry. (B) Confocal images of cytospun E14.5 cardiac cells stained for cTnT (cardiomyocytes), VIMENTIN (non-cardiomyocytes), and DNA, confirming intact nuclear and membrane morphology. Scale bar: 50 µm. (C) Flow cytometric analysis of dissociated E14.5 cardiac cells, identifying ∼35% cTnT+ cardiomyocytes within the nucleated cell population, consistent with the expected composition of the E14.5 murine heart. (D) Agarose gel electrophoresis of MNase-digested chromatin from 500,000 pooled E14.5 cardiac cells (n = 3 hearts) across increasing MNase concentrations (0.5–10 U, 5 min). Lane 1: no-MNase control. (E) Comparative MNase digestion at 3 U for 2.5 min using 250,000 cardiac cells from pooled (n = 3 hearts) or individual (n = 1 heart) E14.5 embryos, demonstrating equivalent nucleosomal profiles and confirming that single embryonic hearts provide sufficient material for chromatin profiling. (F) Workflow schematic for automated adult mouse cardiomyocyte isolation using the gentleMACS Octo Dissociator with Heaters and Heart Perfusion Kit. (G) Representative confocal images of cytospun adult cardiomyocytes stained for ACTN2, MitoSpy Green, and DNA, confirming intact sarcomeric organization, mitochondrial network integrity, and high-purity cardiomyocyte recovery (n = 3). Scale bar: 100 µm. (H) Sequential flow cytometry gating of langendorf-isolated adult cardiomyocytes, showing 96.4% intact cells, 88.2% nucleated (DNA+) events, and 93.4% cTnT+ cardiomyocytes within the nucleated fraction. (I) Representative confocal images of langendorf-isolated and cytospun adult cardiomyocytes stained for ACTN2, cTnT/TNNT2, and DNA, confirming intact sarcomeric organization and high-purity cardiomyocyte recovery (n = 3). Scale bar: 100 µm. (J) Time-course MNase digestion of 250,000 adult cardiomyocyte nuclei (adjusted for ∼85% binucleation) at 3 U across 1–10 min, with clear nucleosomal ladder resolution by 2.5 min. (K) MNase concentration titration of 250,000 adult cardiomyocyte nuclei (0.5–10 U, 5 min), with optimal nucleosomal ladder resolution at 3 U.

Confocal microscopy of cytospun cells confirmed intact nuclear morphology and permitted identification of cTnT-positive cardiomyocytes and VIMENTIN-positive non-cardiomyocyte populations (**Fig. 5B**). Flow cytometric analysis of nucleated cells revealed that approximately 35% of the dissociated cells were cTnT-positive, consistent with the expected cellular composition of the E14.5 murine heart (**Fig. 5C**).

We next performed an MNase titration (0.5–10 U, 5 minutes) using 500,000 cardiac cells pooled from E14.5 embryos (*n* = 3). Agarose gel electrophoresis demonstrated a clear dose-dependent digestion profile, with progressive enrichment of mononucleosomal fragments at higher enzyme concentrations (**Fig. 5D**), mirroring the kinetics previously observed in hiPSCs and hiPSC-CMs. To evaluate whether our protocol could accommodate the limited material available from individual embryonic hearts, we compared MNase digestion of 250,000 cells derived from pooled hearts (3 hearts per tube) versus single hearts (1 heart per tube) at a fixed concentration (3 U, 2.5 minutes). Both conditions produced comparable nucleosomal profiles (**Fig. 5E**), demonstrating that individual E14.5 hearts yield sufficient material for robust chromatin fragmentation without the need for sample pooling. Collectively, these results extend the applicability of our optimized workflow from cultured human pluripotent and differentiated cells to primary murine embryonic tissue, establishing a versatile platform for nucleosomal profiling across both *in vitro* and *in vivo* cardiac models.

### Isolation, enrichment, and MNase-based chromatin digestion of adult mouse cardiomyocytes

Adult cardiomyocytes are among the most technically demanding cell types for chromatin studies. Their large size, rod-shaped morphology, abundant mitochondria, rigid sarcomeric architecture, and predominantly binucleated and polyploid state pose major challenges for dissociation, handling, and enzymatic digestion [17,19,39]. To test the limits of our workflow, we applied it to adult mouse cardiomyocytes isolated using two independent strategies: an automated commercial platform and a previously validated Langendorff-based perfusion method (**Fig. 5F-K**) [56].

We first employed the gentleMACS™ Octo Dissociator combined with the Miltenyi Heart Perfusion Kit, enabling automated tissue digestion, cardiomyocyte enrichment, and recovery of purified cells from whole adult mouse hearts (**Fig. 5F, G**). Confocal microscopy of cytospun cardiomyocytes confirmed that the isolation procedure preserved the characteristic rod-shaped morphology and intact sarcomeric organization, as visualized by α-actinin (ACTN2) and DNA staining. Additional staining with MitoSpy revealed well-organized mitochondrial networks, further confirming the structural integrity of isolated cells (**Fig. 5G**). In parallel, adult cardiomyocytes were independently isolated using the standardized Langendorff perfusion and immunomagnetic purification protocol established by us [56], which has been validated across postnatal ages and yields high-purity cardiomyocyte preparations from individual murine hearts. This allowed us to assess whether our MNase workflow is compatible with cardiomyocytes prepared by an orthogonal isolation strategy. Flow cytometric analysis using sequential gating revealed 96.4% intact cells by scatter profile, 88.2% nucleated (DNA-positive) events, and 93.4% cTnT-positive cardiomyocytes within the nucleated fraction (**Fig. 5H**). Confocal microscopy of cytospun preparations co-stained for ACTN2, cTnT, and DNA corroborated these findings, confirming high-purity cardiomyocyte recovery (**Fig. 5I**).

We assessed MNase digestion kinetics using 250,000 adult cardiomyocyte nuclei, where cell number was adjusted for a calculated ∼85% binucleation rate. A time-course titration at a fixed enzyme concentration (3 U; 1–10 minutes) revealed progressive chromatin fragmentation, with clear nucleosomal ladder resolution emerging by 2.5 minutes (**Fig. 5J**). A complementary enzyme titration (0.5–10 U; 5 minutes) demonstrated dose-dependent enrichment of mono- and dinucleosomal fragments, with 3 U providing optimal ladder resolution (**Fig. 5K**). Notably, adult cardiomyocyte chromatin exhibited digestion kinetics comparable to those observed in hiPSCs, hiPSC-CMs, and embryonic cardiac cells, regardless of the isolation method used, indicating that our standardized conditions are broadly applicable despite the marked differences in cellular architecture, sample processing, cell-type enrichment, and chromatin compaction across these cell types.

## DISCUSSION

Nucleosome positioning and chromatin accessibility are fundamental determinants of gene regulation [11,12,20–22,37,39], yet the technical fragility of MNase-based profiling has confined it to a handful of well-behaved cell types and high-input settings. We address this by establishing a single, systematically optimized MNase-seq workflow that spans cell preparation (i.e., dissociation, enrichment, fresh and fixed), cell type and complexity, cell input scaling, buffer formulation, reverse crosslinking, enzyme concentration, DNA purification, quantitative normalization, sequencing and MNase-seq data analysis (**Fig. 6**). This workflow performs robustly across cell types covering the full range of cell complexity, chromatin biology and technical difficulty: human iPSCs, iPSC-derived cardiomyocytes at successive differentiation stages, primary murine embryonic cardiac tissue, and adult mouse cardiomyocytes. The result is a rapid, scalable MNase framework that lowers both the technical and financial barriers to high-resolution interrogation of nucleosome dynamics in complex biological systems.

**Figure 6.**
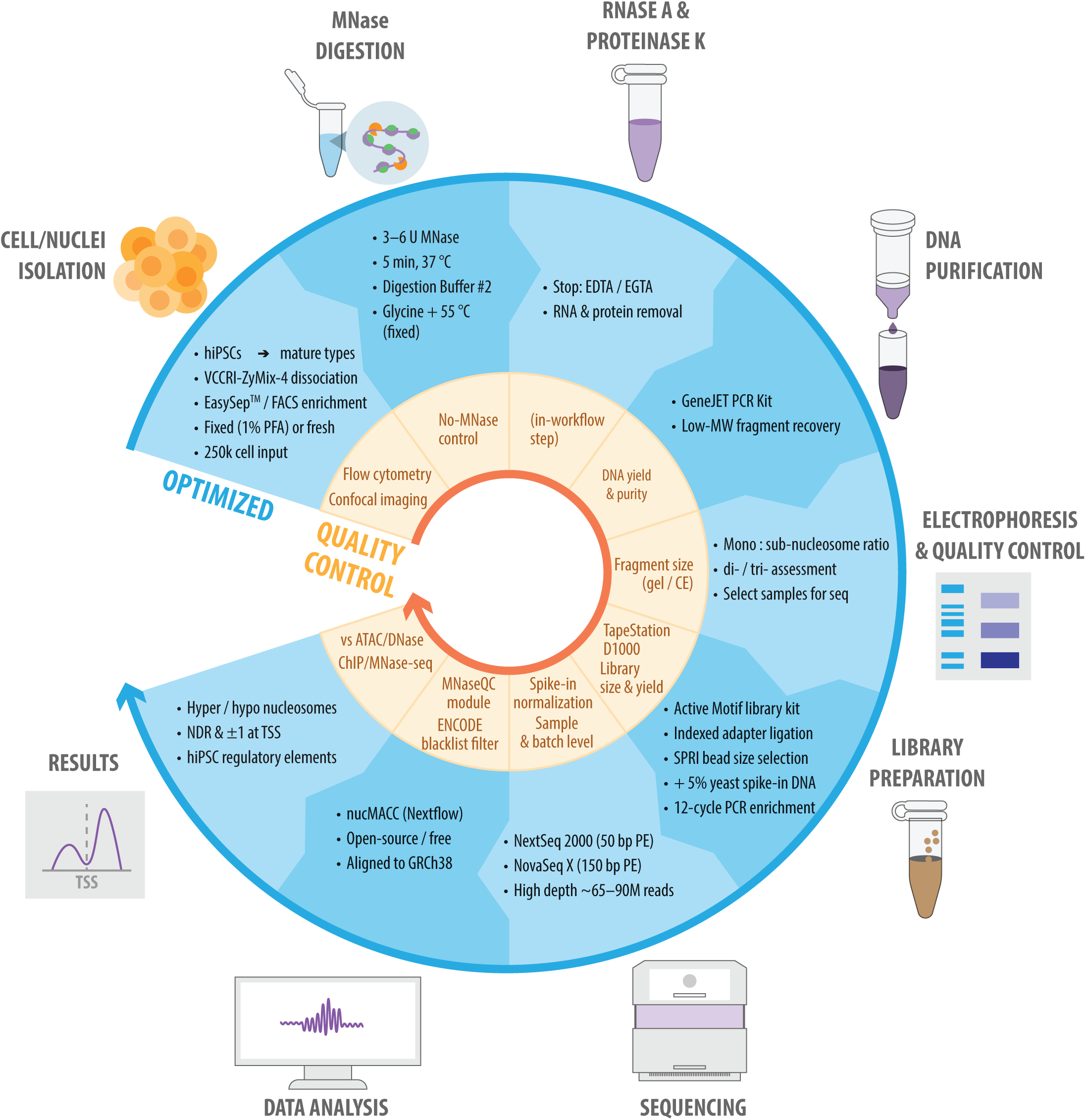
The optimized MNase-seq workflow and its quality-control checkpoints. Circular schematic of the end-to-end framework, read clockwise through nine steps from cell/nuclei isolation to results. The outer ring (blue) summarizes the optimized experimental conditions at each step; the inner ring (orange) shows the corresponding quality-control checkpoint. Steps span cell isolation, MNase digestion and reaction stop (EDTA/EGTA), RNase A/Proteinase K treatments, DNA purification, electrophoresis QC, library preparation with yeast spike-in, next-generation sequencing, nucMACC analysis, and identification of hyper-/hypo-accessible nucleosomes validated against orthogonal ENCODE datasets.

A central finding of this study is that these optimized chromatin digestion conditions can accommodate cell types with markedly different properties without requiring extensive case-by-case re-optimization. This standardization obviates the need to trial an array of MNase protocols that have typically been validated for a single cell type [7,12,17,20,41,57,58]. By benchmarking our workflow across hiPSCs, hiPSC-CMs, embryonic cardiac cells, and adult cardiomyocytes, and demonstrating comparable digestion kinetics across all, we provide evidence that the dominant source of variability in MNase experiments lies not necessarily in intrinsic differences in chromatin compaction between cell types, but rather in upstream sample preparation and the choice of chromatin digestion chemistry. This has direct practical implications by enabling laboratories using multiple models to adopt a single protocol and compare results across model systems with confidence.

Our systematic adaptation and evaluation of buffer composition revealed substantial differences in nucleosomal ladder resolution among six published formulations, with Buffer 2 consistently outperforming all others (**Fig. 6**). While previous studies have typically adopted a single buffer without comparative justification, our side-by-side analysis underscores the extent to which this seemingly routine variable can influence downstream nucleosomal preparations and data quality. Similarly, the marked superiority of the GeneJET PCR Kit over the Monarch Genomic DNA Kit for recovering low-molecular-weight fragments highlights that DNA purification, often treated as an interchangeable step, is in fact a critical determinant of subnucleosomal fragment recovery. This is particularly relevant for MNase-seq applications that rely on the precise quantification of subnucleosomal species, such as *nucMACC*, where incomplete recovery of fragments below 150 bp would bias accessibility scores.

The scalability of our protocol to inputs as low as 100,000 cells without loss of fragment resolution addresses a persistent limitation in the field. Many published MNase-seq workflows require 1–10 × 10^6^ to 1 × 10^8^ cells [11,25,58], restricting them to abundant or pooled material; our profiles remained stable from 10^6^ down to 100,000 cells in both hiPSCs and hiPSC-CMs. Combined with validated fixation and storage, this low-input compatibility opens MNase profiling to rare populations, sorted subsets, limited patient-derived material, and retrospective analysis of biobanked specimens, applications largely inaccessible to existing approaches.

The optimization of PFA fixation and thermal reverse crosslinking for fixed cells fills a persistent gap in the MNase literature. Although fixation is standard in ChIP-seq, its use in MNase workflows has been inconsistent and rarely validated. We find that a brief 10–30 min incubation at 55°C restores enzymatic accessibility, whereas prolonged heating degrades yield, contradicting protocols that employ overnight decrosslinking at elevated temperatures [25]. This matters immediately for scMNase-seq [40] and other fixed-cell applications where fragment integrity is paramount, and the finding that fixed cells tolerate up to four weeks at 4°C provides the temporal flexibility required for multi-condition designs and for clinical settings in which collection and processing cannot be synchronized. The accompanying in-house yeast spike-in addresses the often-ignored problem of quantitative normalization [31,32]: it maintained invariant fragment density and TSS footprinting across all MNase concentrations used in hiPSCs, yet is producible by any laboratory with standard yeast culture, circumventing both the cost and the lot-to-lot variability of commercial reagents.

Applied genome-wide in hiPSCs, the workflow coupled to *nucMACC* resolved concentration dependent nucleosome accessibility and positioning at regulatory elements. Pluripotent cells maintain an open, dynamic chromatin state that underlies broad transcriptional competence [25,58–61], which differentiation remodels through extensive nucleosome repositioning [6,25,59,61]. To our knowledge, this is the first application of a refined MNase-seq protocol with *nucMACC* in hiPSCs to map genome-wide nucleosome and subnucleosome positions, accessibility, and stability across a titration series, establishing a quantitative baseline for the pluripotent epigenome.

Profiling independent ATAC-seq and DNase-seq signal around *nucMACC*-classified mononucleosome centers, hyper-accessible nucleosomes showed the strongest open-chromatin enrichment and hypo-accessible nucleosomes the weakest, with the full set intermediate; this graded relationship was concordant across both orthogonal assays. The same stratification extended to the histone-modification landscape: hyper-accessible nucleosomes were preferentially decorated with active marks (H3K4me3, H3K27ac, and enhancer-proximal H3K4me1), whereas hypo-accessible nucleosomes were enriched for repressive H3K9me3, placing them within constitutive heterochromatic domains. H3K27me3 partitioned mainly with hyper-accessible nucleosomes, co-occurring with active marks at bivalent promoters, with only minor enrichment at hypo-accessible positions. This coherent convergence across accessibility and chromatin-modification dimensions, generated from entirely independent and published datasets, demonstrates that our workflow combined with *nucMACC* does not merely rank MNase signal intensity but resolves biologically meaningful chromatin states embedded in the regulatory architecture of the pluripotent epigenome, and validates our optimized workflow as a quantitative readout of functional chromatin organization.

Identifying VCCRI-ZyMix-4 as the optimal dissociation solution for hiPSC-derived cardiomyocyte cultures and pairing it with EasySep™ magnetic and FACS-based enrichment strategies provide a generalizable template for adapting MNase workflows to mixed cell culture populations (**Fig. 6**). The observation that chromatin digestion kinetics remained relatively consistent between day 8 cardiac progenitors and day 18 cardiomyocytes suggests that the maturation-associated changes in chromatin organization during cardiac differentiation do not alter the efficiency of MNase-mediated chromatin digestion, at least at the level of bulk nucleosomal fragmentation. Whether subtle differences in accessibility at specific loci emerge under these conditions is a question best addressed by MNase-seq and analysis, which our validated pipeline now enables.

The progressive validation from hiPSC-CMs to primary embryonic cardiac tissue and finally to adult mouse cardiomyocytes represents, to our knowledge, a robust cross-model validation of a single MNase workflow reported in the cardiac chromatin field. Each of these models introduced distinct technical challenges. Embryonic hearts required enzymatic dissociation of intact tissue with limited cell yields, yet sufficient material for robust nucleosome profiling was recovered from individual E14.5 hearts without pooling hearts. Adult cardiomyocytes are regarded as among the most difficult cell types to process for cellular and molecular studies due to the specialized isolation methods required, large cell size, highly complex subcellular organization, and rigid sarcomeric structure. Our demonstration that adult cardiomyocyte chromatin digests with kinetics comparable to hiPSCs, using our standardized chromatin digestion method with cardiomyocytes isolated by two independent methods (automated gentleMACS™ dissociation and Langendorff-based perfusion), provides strong evidence for the technical advances of our experimental conditions. The concordance between isolation methods is particularly notable, as it indicates that the MNase digestion step itself is robust to variation in upstream tissue processing—a reassuring finding for studies that may need to integrate data generated across different laboratories or isolation methods.

Looking forward, the unified framework established herein enables quantitative MNase-seq across hiPSC-CM maturation stages, embryonic hearts, and adult cardiomyocytes, opening systematic comparison of nucleosome positioning in cardiac development and disease that has until now been confounded by technical barriers in MNase workflows. Its modular design, with independently validated steps for dissociation, cell enrichment, fixation, reverse crosslinking, chromatin digestion, DNA purification, and sequencing normalization, allows individual components to be adopted within our MNase workflow. By providing detailed optimization data, validated conditions, quality control and a low-cost normalization strategy, we aim to make robust MNase-mediated chromatin profiling reproducible and accessible to a substantially wider community.

## LIMITATIONS OF THE STUDY

This study has two main limitations. First, we validated the workflow by bulk nucleosomal fragmentation across hiPSCs, hiPSC-CMs, mouse embryonic heart cells, and mouse adult cardiomyocytes, but generated genome-wide MNase-seq data only in hiPSCs. Although comparable digestion kinetics are consistent with similar performance across cell types, differences in chromatin compaction and nuclear architecture between pluripotent and post-mitotic states could shift accessibility profiles independently of bulk fragmentation, and formal genome-wide validation in each model remains an important next step. Second, although the workflow accommodates inputs as low as 100,000 cells, further optimization would be required for ultra-low-input or single-nucleus applications, where chromatin digestion and DNA purification must occur on individual nuclei.

## EXPERIMENTAL MODEL AND STUDY PARTICIPANT DETAILS

### Ethics statement and mouse strain

Wild type [WT, Inbred C57BL/6J] (Jackson Laboratory; 000664) mice were bred and housed under pathogen-free conditions in the BioCORE facility (Victor Chang Cardiac Research Institute, Sydney, Australia). Adult female C57BL/6J mice (3-months-old) were maintained on a 12-h light/dark cycle from 7 A.M. to 7 P.M. and had unrestricted access to food and water. All experimental procedures were approved by the Garvan Institute/St. Vincent’s Hospital Animal Experimentation Ethics Committee (No. 19/07, 19/14, 25/16), and performed in strict accordance with the National Health and Medical Research Council (NHMRC) of Australia Guidelines on Animal Experimentation and completed in agreement with The Australian Code of Practice for the Care and Use of Animals for Scientific Purposes (2021). All efforts were made to minimize animal suffering. We used 2-4-month-old WT male and female animals for breeding in our studies.

## METHOD DETAILS

### hiPSC Cell Culture

The male XY hiPSC line PB010.5 (MCRIi010-A (RRID:CVCL_UK91)) with the incorporation of a reporter YFPhGEMININ/mCherryhCDT1 FUCCI cassette into the EEF2 locus, which exhibited no karyotypic abnormalities, was generated by the iPSC Derivation & Gene Editing Facility (Murdoch Children’s Research Institute, Melbourne, Australia). These hiPSCs were split using ReLeSR™ (100-0483, STEMCELL Technologies) and maintained in mTeSR™ Plus (100-0276, STEMCELL Technologies) with 0.5% v/v Penicillin-Streptomycin (15140122, Gibco) under feeder-free conditions on hESC pre-screened Matrigel (354277, Corning). hiPSCs were used between passages 30 to 45. To ensure the quality and pluripotency status of hiPSCs over long-term maintenance, we periodically conducted standard flow cytometric immunophenotyping (Miltenyi Biotec-tested panel 25, Miltenyi Biotec), as well as assessed the growth rate and morphology of hiPSC colonies. hiPSCs were periodically tested for mycoplasma contamination using PCR at the Garvan Institute of Medical Research.

### Optimized MNase-mediated chromatin digestion of native hiPSCs

The optimized MNase digestion workflow developed in this study was applied to undifferentiated FUCCI-reporter hiPSCs (PB010.5) and was the basis for all subsequent adaptations to differentiated and primary cell types. hiPSCs were dissociated to single cells with TrypLE (12604013, Thermo Fisher Scientific), counted on a Countess 3 Automated Cell Counter (Thermo Fisher Scientific), and the required number of unfixed cells (typically 100,000 to 1,000,000) was aliquoted into 1.5 mL Eppendorf tubes. Cells were pelleted at 300 g for 4 min at 4°C and the supernatants gently removed using a P200 pipette. Each pellet was resuspended by flicking the tube, after which 200 μL of MNase digestion Buffer #2 (50 mM NaCl, 10 mM Tris-HCl pH 7.5, 5 mM MgCl2, 1 mM CaCl2, 0.1% (v/v) NP-40, 1 mM β-mercaptoethanol in ultrapure water; adapted from [62]) was added and samples were pipetted up and down five times with a P200 pipette. Cells were permeabilized on a Thermomixer (Eppendorf) at 22°C and 450 RPM for 10 min. Tubes were then removed from the Thermomixer and CaCl2 was added to a final concentration of 2 mM (4 μL of a 100 mM stock per tube) followed by a brief vortex. The Thermomixer temperature was raised to 37°C, and digestion was initiated by adding 2 μL of 1.5 U/μL MNase (EN0181, Thermo Fisher Scientific) per tube to give 3 U per reaction. The 1.5 U/μL working stock was prepared fresh on the day of each experiment by diluting the 300 U/μL MNase stock into MNase Buffer (50 mM Tris-HCl, 5 mM CaCl2, pH 7.9). Tubes were briefly vortexed and incubated at 37°C and 450 RPM for 5 min. Digestion was stopped by addition of 25 μL ice-cold “STOP Solution” (50 mM EDTA, 50 mM EGTA in ultrapure water) directly to each tube while still on the Thermomixer, followed by a brief vortex. RNA was removed by adding 3 μL of 20 mg/mL RNase A (T3018L, New England Biolabs) and incubating at 42°C for 30 min on a heating block. Proteins were digested by adding 12.5 μL of 20% (w/v) SDS together with 2 μL of 20 mg/mL (800 U/mL) Proteinase K Molecular Biology Grade (P8200AA or P8107S, New England Biolabs) and incubating at 55°C for 15 min. Digested DNA was purified using the GeneJET PCR Purification Kit (K0701, Thermo Fisher Scientific) following the manufacturer’s instructions, with elution in 50 μL of 60°C-prewarmed elution buffer. DNA concentration was quantified on a NanoDrop spectrophotometer (NanoDrop® ND-1000 UV-Vis Spectrophotometer, Thermo Fisher Scientific), and fragment size distributions were assessed by agarose gel electrophoresis (2.5% w/v agarose in 1× TAE buffer; Meridian Bioscience Molecular Grade Agarose, cat. #BIO41025), using a 100 bp low-range DNA ladder as a size marker (GWS-DMW-100L, GeneWorks). The ’No-MNase’ control omitted the MNase addition step but was otherwise processed identically.

### Comparison of MNase digestion buffer compositions

To define the optimal digestion buffer for nucleosomal ladder resolution in hiPSCs, six published buffer formulations were evaluated in parallel. FUCCI-reporter hiPSCs were dissociated and counted, and 250,000 cells (No-MNase control), 250,000 cells, or 500,000 cells were aliquoted into 18 1.5 mL Eppendorf tubes (three tubes per buffer). Cells were pelleted at 300 g for 4 min at 4°C, supernatants gently removed, and 200 μL of the corresponding test buffer was added. The buffers compared were: Buffer #1, 10 mM HEPES, 0.5% (v/v) NP-40 in ultrapure water [19]; Buffer #2, as described above [62]; Buffer #3, 10 mM Tris-HCl pH 7.5, 3 mM MgCl2, 0.5% (v/v) NP-40, 10 mM NaCl, supplemented with one EDTA-free Complete Mini protease inhibitor tablet (Roche) [63]; Buffer #4, 20 mM Tris-HCl pH 8.0, 50 mM EDTA, 200 mM NaCl, 1% (w/v) SDS [64]; Buffer #5, 10 mM Tris-HCl pH 7.5, 15 mM NaCl, 60 mM KCl, 3 mM CaCl2 [65]; and Buffer #6, 10 mM Tris-HCl pH 8.0, 1 mM CaCl2 in nuclease-free water [57]. Cells were permeabilised, supplemented with 2 mM CaCl□, and digested with 3 U MNase at 37°C for 5 min. Digestion was stopped by addition of STOP Solution, and DNA was purified using the GeneJET PCR Purification Kit (Thermo Fisher Scientific). Equal volumes of purified DNA (20–25 µL) were loaded into each lane and resolved by agarose gel electrophoresis to compare nucleosomal ladder quality across buffers (Fig. 1B).

### Comparison of DNA purification chemistries

To benchmark DNA purification chemistry, the Monarch Genomic DNA Purification Kit (T3010, New England Biolabs; NEB) was compared with the GeneJET PCR Purification Kit (K0701, Thermo Fisher Scientific) using identical MNase-digested chromatin inputs. FUCCI-reporter hiPSCs (500,000 unfixed cells per condition) were aliquoted across six tubes corresponding to a digestion gradient (No-MNase, 0.5 U, 1 U, 3 U, 6 U, 10 U) and processed through the optimized digestion workflow with a 10 min incubation at 37°C. After Proteinase K treatment, each lysate was split into two equal aliquots (∼125 μL each) and processed in parallel using the two kits according to manufacturers’ instructions. For Monarch, samples were mixed with 400 μL gDNA binding buffer, transferred to gDNA columns, centrifuged (3 min at 1,000 g, then 1 min at 15,000 g), and washed twice with 500 μL gDNA wash buffer at 15,000 g. DNA was eluted in 50 μL of 60°C-prewarmed gDNA elution buffer. For GeneJET, samples were mixed with an equal volume of binding buffer, supplemented with 10 μL of 3 M sodium acetate (pH 5.2) and an equal volume of 100% isopropanol, transferred to GeneJET columns, centrifuged at 15,000 g for 1 min, washed once with 700 μL wash buffer, and eluted in 50 μL of 60°C-prewarmed elution buffer. Eluates (20 µL) were resolved by agarose gel electrophoresis to assess recovery of mononucleosomal and subnucleosomal fractions.

### MNase enzyme titration and time-course analysis

Dose-response and kinetic profiles of MNase digestion were characterized in unfixed hiPSCs using the optimized workflow. For the enzyme titration, 500,000 cells per tube were distributed into nine tubes corresponding to No-MNase, 0.5, 1, 3, 6, 10, 15, 30 and 50 U MNase. Working dilutions were prepared in MNase Buffer: a 25 U/μL stock was first generated by diluting 2 μL of 300 U/μL MNase into 22 μL MNase Buffer, and serial dilutions yielded 15, 7.5, 5, 3, 1.5, 0.5 and 0.25 U/μL working solutions, of which 2 μL was added per tube to deliver the indicated unit doses. Reactions were incubated at 37°C and 450 RPM for 5 min and processed as in the optimized workflow. For the time-course, 500,000 cells per tube were distributed into 13 tubes (No-MNase, plus 1 U or 3 U at 1, 2.5, 5, 10, 15, and 30 min). MNase additions were staggered in descending order of time point so that all reactions were stopped simultaneously at the 30 min endpoint with cold STOP Solution.

### Cell input scalability of MNase digestion

To define the lower limit of cell input compatible with the optimized workflow, hiPSCs were aliquoted at 100,000, 250,000, 500,000 or 1,000,000 cells per tube, in duplicate (one set for 5 min and one set for 15 min digestion), with an additional 100,000-cell No-MNase control per set. Cells were processed through the optimized workflow with 3 U MNase at 37°C and 450 RPM. The 5 min set was stopped at 5 min and the 15 min set at 15 min by addition of cold STOP Solution. RNase A, Proteinase K, and GeneJET-based purification proceeded as described, and fragment distributions were resolved by agarose gel electrophoresis.

### PFA fixation and storage of hiPSCs for MNase digestion

For fixed-cell MNase analyses, freshly dissociated FUCCI-reporter hiPSCs were resuspended in PBS containing 1% (v/v) paraformaldehyde (PFA, prepared by diluting 4% PFA stock in PBS) and incubated at room temperature or on ice for 10 min. Crosslinking was quenched by addition of glycine to 125 mM final concentration (52.5 μL of a 2.5 M stock per 1 mL sample) followed by 5 min incubation. Fixed cells were washed in PBS or 1% (w/v) BSA in PBS, pelleted at 500 g for 5 min, and stored at 4°C in PBS containing 0.02% (w/v) sodium azide. To compare freshly fixed and stored material, FUCCI-reporter hiPSCs were either dissociated and fixed two weeks before the experiment or freshly dissociated on the day of digestion; freshly dissociated cells were further split into a fixed-and-quenched and an unfixed-and-unquenched arms. Cells from each condition were aliquoted at 250,000 or 500,000 cells per tube and processed in parallel using the optimized digestion workflow with 3 U MNase for 5 min at 37°C.

### Optimization of PFA concentration and synergy with thermal decrosslinking

To define the PFA concentration window compatible with downstream MNase digestion, and to test whether thermal decrosslinking rescues digestion at higher fixative concentrations, two parallel experiments were performed using freshly dissociated FUCCI-reporter hiPSCs. In both arms, 250,000 cells were aliquoted into six tubes and resuspended in PBS containing 0% (no-PFA control), 0% (no-PFA, no-quench control), 0.1%, 0.5%, 1% or 2% (v/v) PFA in a final volume of 1 mL, and fixed on ice for 10 min. In the first arm (chemical quenching only), tubes were quenched with 125 mM glycine for 5 min on ice, washed in 1% (w/v) BSA in PBS, pelleted at 500 g for 5 min, resuspended in 200 μL Digestion Buffer #2, and digested with 3 U MNase for 2.5 min at 37°C and 450 RPM. In the second arm (chemical quenching plus rapid thermal decrosslinking), cells were quenched with glycine, pelleted, resuspended in 1 mL PBS containing 125 mM glycine, and incubated at 55°C for 30 min on a heating block prior to MNase digestion under identical conditions. The two arms were processed side-by-side to compare the resolution of nucleosomal ladders obtained with and without thermal decrosslinking across the PFA concentration range.

### Optimization of thermal decrosslinking

To establish a decrosslinking regime that restores enzymatic accessibility without compromising DNA integrity, FUCCI-reporter hiPSCs were dissociated, fixed in 1% PFA for 10 min on ice, and quenched with 125 mM glycine for 5 min. After washing in PBS and centrifugation at 500 g for 5 min, cells were counted and 400,000 cells were aliquoted into Eppendorf tubes corresponding to four heating durations (no glycine and 0 min heating; 30 min; 60 min; or overnight, 18 h). An additional 200,000-cell aliquot was retained as a No-MNase control. Tubes were made up to 1 mL in FACS buffer (PBS, 2% (v/v) FBS, 2 mM EDTA pH 7.9), 125 mM glycine was added at the indicated time points, and tubes were incubated at 55°C on a heating block for the corresponding duration. The fine-scale decrosslinking time series used a comparable design with 250,000 cells per tube and five conditions in duplicate (no heating, 10 min, 30 min, 45 min, or 60 min at 55°C). After heating, samples were processed through the optimized digestion workflow using 3 U MNase for 2.5 min at 37°C.

### Yeast culture for spike-in DNA generation

*Saccharomyces cerevisiae* strain BY4742 was used as the source of mononucleosomal spike-in DNA. A stab from a glycerol stock was inoculated into 2 mL of filter-sterilized YEPD medium (1% (w/v) yeast extract, 2% (w/v) Bacto peptone, 2% (w/v) dextrose) in a 15 mL tube and incubated at 30°C for 4 h without shaking, followed by 22 h at 30°C with shaking at 220 RPM, and stored at room temperature. An overnight starter culture was prepared by transferring 25 μL of the stab culture into 4 mL of fresh YEPD and incubating at 30°C and 220 RPM for 17 h, followed by storage at room temperature. To generate log-phase cultures for digestion, 10 μL of the overnight starter was inoculated into 4 mL of YEPD and grown at 30°C and 220 RPM for 16 h; on the following morning, 1 mL of this culture was transferred into 25 mL of fresh YEPD in a 50 mL tube and OD600 was monitored at regular intervals. Cells in mid-log phase (OD600 = 0.4 to 0.9; harvest typically at OD600 ≈ 0.9) were pelleted at 2,500 g for 6 min. Most supernatant was removed and pellets were resuspended in 750 μL PBS, after which 4% PFA stock was added to a final concentration of 1% and cells were fixed at room temperature for 10 min. Crosslinking was quenched with 125 mM glycine, and cells were transferred to 1.5 mL Eppendorf tubes, pelleted at 20,000 g for 90 s, and stored at -80°C until use.

### MNase digestion of yeast chromatin and custom-made spike-in DNA preparation

Frozen BY4742 pellets were resuspended in 1 mL Complete Spheroplast Buffer (1 M sorbitol, 50 mM Tris pH 7.5, 5 mM β-mercaptoethanol, 200 U/mL Zymolyase (E1004, Zymo Research)) and incubated at 25°C and 600 RPM for 30 min on a Thermomixer. Spheroplasts were pelleted at 20,000 g for 30 s, resuspended in 200 μL Digestion Buffer (Buffer #2 above), and supplemented with 4 μL of 100 mM CaCl2 (final 2 mM) and 60 U MNase. Digestion proceeded at 37°C and 600 RPM for 10 min and was stopped with 25 μL STOP Buffer (50 mM EDTA, 50 mM EGTA). Samples were treated with 6 μL of 20 mg/mL RNase A at 42°C for 30 min, followed by 25 μL of 20% (w/v) SDS and 10 μL of 20 mg/mL Proteinase K at 60°C for 45 min. DNA was purified using the GeneJET PCR Purification Kit, quantified on a NanoDrop, and stored at -20°C. Yeast spike-in DNA was added to each hiPSC MNase-derived DNA sample at a fixed ratio (0.5 ng yeast DNA per 10 ng hiPSC DNA, equivalent to ∼5% of total) immediately prior to library preparation.

### Sequencing library preparation and MNase-seq

NGS libraries were generated from yeast-spiked MNase-digested hiPSC DNA using the Active Motif DNA Library Kit (DNA Library Prep Kit for Illumina, 53220) according to the manufacturer’s instructions. Briefly, 10 ng of fragmented hiPSC DNA combined with 0.5 ng of BY4742 spike-in DNA was diluted in 10 mM Tris-HCl to a final volume of 39 μL, mixed with 10 μL End Repair buffer and 1 μL End Repair Enzyme Mix, and incubated in a thermal cycler at 12°C for 15 min, 37°C for 15 min, 72°C for 30 min, and held at 4°C. Diluted Illumina adapters (2.5 μL), 15 μL PEG and 1.25 μL T4 DNA ligase were then added, and ligation proceeded at 25°C for 15 min without a heated lid. Adapter-ligated DNA was purified with 55 μL of room-temperature SPRI beads, washed twice with 200 μL of 80% (v/v) ethanol on a magnetic stand (DynaMag-2, Thermo Fisher, 12321D), and eluted in 22 μL TT buffer. Adapter-ligated libraries (20 μL) were enriched by PCR using 2.5 μL each of AM i5- and i7-XXX primers and 25 μL Q5 Polymerase NGS master mix with the program: 98°C for 2 min; 12 cycles of 98°C for 10 s and 65°C for 75 s; 65°C for 2 min; hold at 4°C. PCR-enriched libraries were size-selected with 37.5 μL SPRI beads, washed twice with 200 μL of 80% (v/v) ethanol, and eluted in 17 μL of 37°C-prewarmed TT buffer. 15 μL of each library was transferred to a fresh 200 μL PCR tube and stored at -20°C until sequencing. Library quality and quantity was assessed by capillary electrophoresis with the TapeStation (Agilent Technologies) automated electrophoresis system with TapeStation D1000 ScreenTape and reagents (Agilent Technologies), and libraries were sequenced on Illumina NextSeq (50-bp paired-end) or NovaSeq-X (150-bp paired-end) platforms by the Australian Genome Research Facility (AGRF). Reads were processed and analyzed using the *nucMACC* pipeline (see below).

### MNase-seq reads processing, input data, and *nucMACC* implementation

Input data consisted of paired-end FASTQ files generated from MNase digestion experiments. Sequencing reads were supplied to the pipeline via a structured sample sheet specifying sample identifiers and file paths to forward and reverse reads.

The *nucMACC* workflow (version 3.1) was executed using *Nextflow* (version 25.10.4), with analysis modules configured for MNase quality control (MNaseQC) and nucleosome profiling (nucMACC) using the --analysis parameter. Pipeline execution included specification of the sample sheet (--csvInput), output directory (--outdir), GRCh38 reference genome and index (--genome, --genomeIdx), genome size (--genomeSize), transcription start site annotation file (--TSS), and a blacklist of 636 problematic genomic regions derived from the ENCODE hg38 Blacklist v2 (--blacklist) [66]. Filtered BAM outputs were generated using the --publishBamFlt parameter. The pipeline was deployed across multiple computational environments, including Azure cloud-based virtual machines and high-performance computing infrastructure (NCI Gadi), with configuration adjustments implemented to accommodate file system and execution constraints.

### Processing of low-input and high-input datasets

Low-input MNase-seq data (5,930,979–8,408,494 reads) were generated using an Illumina *NextSeq 2000* platform with 50 bp paired-end sequencing and processed using the *nucMACC* Nextflow pipeline. The quality control module was executed successfully, producing eight output directories. Fragment size statistics generated in the 06_FragmentStatistics module indicated an increase in sub-nucleosomal fragments with increasing MNase concentration. However, the nucleosome profiling module failed during execution of the get_nucMACC_scores.R script. Investigation revealed the presence of a hard-coded read depth filter within the script, suggesting that insufficient sequencing depth contributed to the failure. Consequently, low-input datasets were not pursued further for downstream analysis.

To address this limitation, the same MNase libraries were re-sequenced on an Illumina *NovaSeq X* platform (XLEAP-SBS chemistry, patterned flow cell) to generate higher-depth 150 bp paired-end datasets (65,362,382–87,859,308 reads). Quality control analysis completed successfully, again producing eight output directories. In contrast to the low-input dataset, fragment statistics reported predominantly mono-nucleosomal fragments with no detectable sub-nucleosomal population.

To investigate this discrepancy, sequencing reads were cropped to 50 bp prior to analysis, under which conditions both mono-nucleosomal and sub-nucleosomal fragments were detected. Comparative analysis of sub-nucleosomal alignments derived from cropped 50 bp reads and mono-nucleosomal alignments from full-length 150 bp reads demonstrated concordant genomic positioning, indicating that the absence of sub-nucleosomes in the 150 bp dataset was unlikely to reflect a biological difference. Instead, the discrepancy was attributed to adapter contamination: when MNase-digested fragments are shorter than the 150 bp read length—as is the case for sub-nucleosomal fragments (∼80–150 bp)—adapter sequence is incorporated into the 3’ end of each read. In the absence of trimming, this contaminant sequence causes erroneous insert size estimation during alignment, misclassifying short fragments into larger size categories. To address this, FASTQ files were processed using *Trimmomatic* (version 0.40) with the following parameters:

ILLUMINACLIP:adapters/TruSeq3-PE.fa:2:30:10 LEADING:3 TRAILING:3 SLIDINGWINDOW:4:15 MINLEN:33

Following trimming, *nucMACC* analysis failed to recover sub-nucleosomal fragments in certain samples (notably 10 U MNase conditions), attributable to reduced read depth following removal of orphaned reads during trimming. To retain these reads, a modified trimming approach was implemented using Trimmomatic with parameters configured to preserve orphan reads:

ILLUMINACLIP:adapters/TruSeq3-PE.fa:2:30:10:1:true

Retention of orphan reads resulted in failure of the *MNaseQC* module, with errors arising in the *GenerateTxtFragCounts.R* script. Inspection revealed that this script assumes a fixed log file structure and is hard-coded to skip the first four lines before searching for a specific text string. Warning messages associated with orphan reads disrupted this expected format. The script was modified to accommodate variable log file structures, enabling successful completion of the MNaseQC module. Despite these corrections, the nucMACC module continued to fail during execution of the get_nucMACC_scores.R script. Further inspection identified an internal read depth filtering threshold [original value: 31] that resulted in exclusion of all candidate regions. We relaxed this threshold to 14 which enabled recovery of mono-nucleosomal outputs in our high-depth datasets. We also applied and evaluated other thresholds, including 8 and 11.

### Genomic feature overlap and enrichment analysis

The genomic distribution of all mononucleosomes, hyperaccessible nucleosomes, and hypoaccessible nucleosomes was assessed by intersecting these regions with genomic annotations from GENCODE v48 (hg38). Overlap analyses were performed using BEDTools v2.31.1 [67], specifically the intersect function with the parameters -wao -f 0.40, requiring a minimum overlap of 40% between the nucleosome region and the annotated genomic feature. The resulting overlaps were used to quantify the enrichment of nucleosome classes across genomic features, including promoters, exons, introns, untranslated regions (UTRs), and intergenic regions. Feature categories were not mutually exclusive: a nucleosome meeting the 40% overlap threshold for more than one feature was counted in each, so class proportions may sum to more than 100%.

### ENCODE histone modification and chromatin accessibility enrichment analysis

To investigate the chromatin features associated with regions displaying hyperaccessible or hypoaccessible nucleosomes, publicly available chromatin accessibility and histone modification datasets were obtained from the ENCODE project (https://www.encodeproject.org/) for the independent human induced pluripotent stem cell line GM23338. The following datasets were used: ATAC-seq (ENCFF234AYB; signal p-value track), DNase-seq (ENCFF241UWA and ENCFF395HEQ; read-depth normalised signal tracks), and Mint-ChIP-seq foldchange over control signal tracks for the histone modifications H3K27ac (ENCFF234VLU), H3K4me3 (ENCFF521ANU), H3K4me1 (ENCFF121RRE), H3K27me3 (ENCFF846RZH), and H3K9me3 (ENCFF466TTA).

Signal enrichment around hyperaccessible and hypoaccessible nucleosome regions was quantified using deepTools (v3.5.2) [68]. Briefly, signal matrices were generated using the *computeMatrix* function in reference-point mode, centered on the regions of interest and extending (±1 kb) upstream and downstream from each region’s center. Enrichment profiles were visualized using the deepTools *plotHeatmap* and *plotProfile* functions. These analyses were used to compare chromatin accessibility and active and repressive histone modifications across the identified nucleosome classes.

### Processing of public MNase-seq data and correlation assessment

FASTQ files were downloaded using the SRA Toolkit (3.0.10) from BioProject PRJNA254322, SRA study SRP044041, specifically runs SRR1781837 and SRR1781838, which correspond to MNase-seq data from human induced pluripotent stem cells [25]. Adapter trimming and length filtering were performed with Cutadapt v2.4 [69], removing Illumina Nextera adapter sequences (-a CTGTCTCTTATACACATCT -A CTGTCTCTTATACACATCT) and discarding reads shorter than 20 nt (-m 20).

Trimmed reads were mapped to the human hg38 reference genome using Bowtie 2 [70] with the parameters --very-sensitive --fr --no-mixed --no-discordant --dovetail. PCR duplicates were removed using the Picard *MarkDuplicates* function with default settings (http://broadinstitute.github.io/picard/). SAMtools (v1.8) [71] was then used to remove mitochondrial reads and reads with mapping quality <30. BigWig files were generated using the *bamCoverage* function from deepTools with a bin size of 10 bp.

To assess the correlation between our high-depth MNase-seq data and the published West et al. (2014) datasets SRR1781837 and SRR1781838, we used multiBigwigSummary (bins mode) from deepTools, excluding blacklisted regions using ENCFF356LFX.bed. Pairwise correlations were visualised using plotCorrelation with Pearson correlation, heatmap output, and the --skipZeros option.

### Fixation protocol for MNase-mediated chromatin digestion and flow cytometry of hiPSCs

For fixing hiPSCs for flow cytometry, cells in 6-well plates were processed as follows. The media was removed, and each well was washed once with 1 mL of PBS. Then, 1 mL of TrypLE was added to each well, and the plates were incubated for 5-7 minutes at 37°C. The cells were pipetted up and down a few times per well to obtain a single-cell suspension, which was corroborated using a microscope. Next, 1 mL of cold FACS Buffer (PBS 1×, 2% FBS v/v, 2 mM EDTA pH 7.9) was added to each well to inactivate the TrypLE. The cells from each well were transferred to separate 2 mL tubes and spun at 500 g for 5 minutes (300 g for 3 minutes for hiPSCs). The supernatant was removed, and the pellets were resuspended by flicking the tubes. Then, 1 mL of FACS Buffer was added to each well. The tubes were checked for clumps by looking at them against a light source, and any clumps present were removed by flicking the tubes. 4% PFA was added to a final concentration of 1% in each tube, and the cells were fixed for 10 minutes at room temperature. After fixation, the cells were washed with abundant PBS, and the tubes were spun at 400 g for 4 minutes (300 g for 3 minutes for hiPSCs). The supernatant was removed, and the pellets were resuspended by flicking. Finally, 1-2 mL of PBS with 0.02% Sodium Azide was added to each tube, and the samples were stored at 4°C. The volumes were adapted when using smaller well plates.

### Cytospin of cells in suspension

After mounting CytoSep™ Cytology Funnels (base holder and fluid chamber) for the Sakura Cyto-Tek® Cytocentrifuge (Model 4323, Sakura) and a white filter for the Sakura cytology funnel (M963FW, Simport), PFA-fixed cells in suspension were centrifuged (2,000 RPM, 3 min) onto Epredia™ SuperFrost Plus™ Adhesion slides (12312148, Thermo Fisher Scientific). Following the creation of a hydrophobic barrier with a PAP pen (ab2601, Abcam), the cells were ready for staining. We typically used 0.1 to 0.2 mL of cells in suspension.

### EasySep magnetic enrichment of hiPSC-derived cardiomyocytes

Cardiomyocyte enrichment from hiPSC-CM cultures was performed using the EasySep Human PSC-Derived Cardiomyocyte Enrichment Kit (17893, STEMCELL Technologies) with optimizations to minimize non-specific cell loss. After dissociation with the in-house Cardiomyocyte Dissociation Buffer (described below in *hiPSC-Derived Cardiomyocyte In-house Dissociation Protocol*), cells were pelleted at 300 g for 4 min and resuspended in 1 mL FACS buffer. Cells were counted and adjusted to no more than 25 × 10□ cells/mL; samples containing fewer than 2.5 × 10□ total cells were further pelleted and resuspended in 0.1 mL FACS buffer. Cells were transferred to 5 mL polystyrene round-bottom tubes (352054, Corning), and 50 μL of EasySep Enrichment Cocktail was added per mL of sample. Tubes were flicked to mix and incubated at room temperature for 5 min. RapidSpheres were vortexed for 30 s to ensure even dispersion, and 100 μL was added per mL of sample, followed by a further 5 min incubation at room temperature. FACS buffer was added to bring the final volume to 2.5 mL, and the cell suspension was mixed by gentle pipetting (two to three times). Tubes were placed without lids in the EasySep magnet and incubated at room temperature for 5 min. The cardiomyocyte-enriched fraction was collected by inverting the magnet and tube together in a single smooth motion into a new labeled 5 mL polystyrene round-bottom tube.

### MNase digestion of hiPSC-derived cardiomyocytes at multiple differentiation stages

To assess MNase digestion across cardiomyocyte maturation, hiPSC-CMs were harvested at days 8, 12 and 18 of directed differentiation and processed independently using a common protocol. Cells were dissociated, pelleted at 300 g for 4 min, resuspended in 1 mL FACS buffer and counted. Cardiomyocytes were enriched using the EasySep workflow described above, with the exception of day-8 cultures, where enrichment was omitted because of significant cell losses observed at this early differentiation stage. Enriched (or unenriched) cells were aliquoted into four tubes per stage corresponding to a No-MNase control (100,000 cells), 100,000, 250,000 and 500,000 cells. Tubes were pelleted at 300 g for 4 min at 4°C, supernatants removed, pellets resuspended in 200 μL Digestion Buffer #2, and processed through the optimized workflow using 3 U MNase for 5 min at 37°C and 450 RPM. DNA was purified with the GeneJET PCR Purification Kit and resolved by agarose gel electrophoresis to evaluate ladder resolution across stages and inputs.

### MNase enzyme titration in day 8 and day 18 hiPSC-derived cardiomyocytes

Dose-response analyses in differentiated cardiomyocytes were performed at day 8 (without enrichment) and day 18 (with EasySep enrichment) of directed differentiation. For each stage, 500,000 cells were aliquoted into six tubes (No-MNase, 0.5 U, 1 U, 3 U, 6 U and 10 U). Working dilutions of MNase were prepared by diluting 2 μL of the 300 U/μL stock into 78 μL MNase Buffer to give a 7.5 U/μL solution, with serial dilution to 5, 3, 1.5, 0.5 and 0.25 U/μL. 2 μL of the appropriate dilution was added per tube. Tubes were incubated at 37°C and 450 RPM for 5 min and processed through the standard digestion workflow.

### FACS-based purification and MNase digestion of fixed Day 12 hiPSC-derived cardiomyocytes

To validate the fixed-cell MNase workflow in differentiated cardiomyocytes, day-12 hiPSC-CMs were dissociated, fixed in 1% PFA for 10 min, washed in excess PBS, pelleted, and stored in 2 mL PBS at 4°C. Several days later, cells were pelleted at 500 g for 5 min, resuspended in 1 mL of in-house permeabilization/wash buffer, and permeabilized at room temperature for 15 min. Cells were stained overnight at 4°C in 200 μL of permeabilization/wash buffer containing 1:200 PE Mouse Anti-Cardiac Troponin T (cTnT (PE), BD Biosciences #564767, clone 13-11, recognises TNNT2), and 1:5,000 Vybrant DyeCycle Violet (V35003, Thermo Fisher Scientific). cTnT-positive cardiomyocytes were sorted into three aliquots of 250,000 cells each on a BD FACSAria III Cell Sorter and stored at 4°C. Two of the three sorted aliquots were quenched with 125 mM glycine; the third was processed unquenched. All three aliquots were decrosslinked at 55°C for 30 min, pelleted at 500 g for 5 min, resuspended in 200 μL Digestion Buffer #2, and processed through the optimized workflow using 3 U MNase for 2.5 min at 37°C.

### Timed mating and dissection of the mouse embryo

Timed mating was performed between wild-type (WT) male and female mice, with pregnancy confirmed by the detection of a copulation plug. Pregnant females were sacrificed as previously described, and E14.5 embryos were collected in glass Petri dishes containing ice-cold 1X PBS. The embryos were briefly rinsed in ice-cold 1X PBS to remove any residual blood.

### Embryonic heart cell isolation and fixation

Wild-type (WT) E14.5 embryonic hearts from both male and female mice were isolated by surgical removal and cleaned of surrounding tissue. The hearts were then cleared of blood using cold PBS, followed by a single FACS buffer wash. Each heart was dissociated into single cells with the *Cardiomyocyte Dissociation Buffer* (30-40 minutes at 37°C) [54]. The dissociated cells were collected in 2 mL Eppendorf tubes, filtered through a pluriStrainer Mini 100 µm (43-10100-50, pluriSelect) into a new tube, and fixed in ice-cold 1% PFA (in FACS buffer) for 10 minutes. Antibody staining was subsequently performed on 250,000 total cells as previously described.

### MNase digestion of E14.5 embryonic cardiac cells

E14.5 embryonic cardiac cells were dissociated and fixed in 1% PFA as described in *’Embryonic Heart Cell Isolation and Fixation’* above. For the enzyme titration, 500,000 dissociated cells pooled from three E14.5 embryos were aliquoted into six tubes (No-MNase, 0.5, 1, 3, 6 and 10 U). Tubes were made up to 1 mL with PBS and decrosslinked at 55°C for 10 min on a heating block. Cells were pelleted at 500 g for 5 min at 4°C, resuspended in 200 μL Digestion Buffer #2, and digested at 37°C for 5 min using the MNase dilution series described above. To compare pooled and single-heart inputs, 250,000 cells per tube were prepared from either three pooled E14.5 hearts or single E14.5 hearts (two tubes per condition), decrosslinked, and digested with 3 U MNase for 2.5 min at 37°C. All samples were processed through RNase A, Proteinase K, and GeneJET-based purification as described in the optimized workflow.

### Flow cytometry combined with antibody staining of cells

For flow cytometric analysis of cells, the required number of fixed cells was transferred into 1.5 mL Eppendorf tubes. We typically used between 200,000 to 1,000,000 cells. Each tube received 1 mL of 1% BSA in PBS, followed by centrifugation at 500 g for 5 minutes. The supernatants were discarded, and the pellets were resuspended by flicking. Subsequently, 500 µL of 1X in-house permeabilization/wash buffer (0.2% (w/v) saponin containing either 4% FBS (v/v) or 2% NCS (v/v), 1% (w/v) BSA and 0.02% (v/v) Sodium Azide in PBS) was added to each tube, and the samples were incubated at room temperature (RT) for 10 minutes. After incubation, the tubes were centrifuged again at 500 g for 5 minutes. The supernatants were discarded, leaving 50 µL in each tube, and the pellets were resuspended by flicking. 100 µL of primary or conjugated antibodies, prepared in 1X permeabilization/wash buffer, was added to the relevant tubes, and the samples were incubated overnight at 4°C. Antibodies used in this study included anti-cardiac troponin T (BD Pharmingen™ Alexa Fluor® 647 Mouse Anti-Cardiac Troponin T, BD Pharmingen™ PE Mouse Anti-Cardiac Troponin T; or unconjugated anti-cTnT, 1:200, MA5-12960, Thermo Fisher Scientific), Anti-α-Actinin (Sarcomeric) (clone REA402, Vio R667 conjugate, Miltenyi Biotec, REAfinity recombinant antibody), anti-VIMENTIN (1:100, D21H3, XP® Rabbit mAb, Alexa Fluor® 647 Conjugate, Cell Signaling Technology #9856). The next day, the tubes were washed once with 1 mL of 1% BSA in PBS and once with 0.5 mL of 1X perm/wash buffer, followed by centrifugation at 500 g for 5 minutes. The supernatants were discarded, leaving 50 µL in each tube, and the pellets were resuspended by flicking. When using secondary antibodies, 50 µL of them, prepared in 1X permeabilization/wash buffer, was added to the relevant tubes, and the samples were incubated at RT for 60-120 minutes, covered. After this incubation, the tubes were washed once with 1 mL of 1% BSA in PBS and once with 0.5 mL of 1X perm/wash buffer. The supernatants were discarded, leaving approximately 100 µL in each tube, and the pellets were resuspended by flicking. A 1:2500 dilution of Vybrant™ DyeCycle™ Violet (V35003, 5 mM) in 1X perm/wash buffer was prepared, and 100 µL was added to each tube for a final concentration of 1:5000. The tubes were covered and taken for analysis by flow cytometry.

### hiPSC-derived ventricular cardiomyocyte in-house differentiation protocol

On Day -2, hiPSCs were dissociated using pre-warmed TrypLE (TrypLE Express Enzyme (1×), no phenol red, Cat. No. 12604013, Thermo Fisher Scientific, USA) for 7 min and seeded at a density of 200,000 cells/cm² in 24 or 48-well plates using mTeSR medium supplemented with 5 µM Y-27632. On Day -1, the media in the wells was replaced with fresh mTeSR. On Day 0, when the wells were 70-90% confluent, they were washed once with PBS and 6 µM CHIR99021 in RPMI+B27(-ins)+Glutamax was added. Approximately 24 hours later, on Day 1, the wells were washed once with PBS, and RPMI+B27(-ins)+Glutamax with 1 µM CHIR99021 was added. On Day 3/4, the wells were washed once with PBS and RPMI+B27(-ins)+Glutamax supplemented with 200 µM L-Ascorbic acid, 5 µM IWP-2, and 5 µM XAV939 was added. 48 hours after Day 3/4, on Day 5/6, the wells were washed once with PBS and RPMI+B27(-ins)+Glutamax supplemented with 200 µM L-Ascorbic acid added. On Day 7, the media was replaced with RPMI+B27+Glutamax supplemented with 200 µM L-Ascorbic acid. From Day 9 onwards, the media was replaced every 2-3 days with RPMI+B27+Glutamax supplemented with 200 µM L-Ascorbic acid.

### Comparison of dissociation solutions for hiPSC-derived cardiomyocytes and viability assessment

To define optimal dissociation conditions for hiPSC-CMs, four enzymatic dissociation cocktails were tested in parallel across multiple stages of directed cardiac differentiation. The four cocktails used were: 1X Trypsin/EDTA (0.05% Trypsin/0.02% EDTA) (Thermo Fisher Scientific, 15400054) made up in DMEM/F12 (Thermo Fisher, 11320033); Stemdiff cardiomyocyte dissociation kit (Stemcell Technologies, 5025); TrypLE+Collagenase IV (1X TrypLE (Thermo Fisher Scientific, 12604013) + 0.5 mg/mL Collagenase IV (Cat. #LS004188, Worthington); VCCRI-ZyMix-4 (12.5 U/mL Papain (Cat. #10108014001, Sigma Aldrich), 0.5 mM Cysteine, 132.5 U/mL Collagenase II (Cat. #LS004176, Worthington), 0.5% BSA (Cat. #A7906, Sigma), 0.025 mg/mL DNase I (Cat. #10104159001, Sigma Aldrich), and 0.1 U/mL Dispase (Cat. #07913, STEMCELL), all made up in DMEM (+L-Glutamine, +Sodium Pyruvate, +4.5 g/L Glucose). VCCRI-ZyMix-4 corresponds to the optimized in-house Cardiomyocyte Dissociation Buffer described in *hiPSC-Derived Cardiomyocyte In-house Dissociation Protocol*] [54]. For each comparison, sister wells of differentiating hiPSC-CMs in 48-well plates were washed with 0.5 mL/well PBS and incubated with 250 µL/well of the respective room-temperature dissociation cocktail for 30-40 min at 37°C with gentle tapping every 15 min. Day 20-30 hiPSC-CMs needed 40 min at 37°C. Following digestion, cells were quenched with 500 µL FACS buffer per well, gently pipetted up and down 20 times with a P1000 pipette to achieve single-cell status, transferred to 15 mL Falcon tubes, washed with excess FACS buffer, and pelleted at 300 g for 4 min at room temperature. Cell viability was assessed immediately on freshly dissociated, unfixed cells by co-staining with 250 nM Calcein-AM (C3100MP, Thermo Fisher Scientific) and 1 µg/mL propidium iodide (PI; P4170, Sigma Aldrich) in PBS for 10 min at 37°C protected from light, followed by flow cytometric analysis on a BD LSRFortessa™ X-20 Cell Analyzer using 488 nm excitation (Calcein-AM, FITC channel) and 561 nm excitation (PI, PE-Texas Red channel). Viability was quantified as the proportion of Calcein-AM-positive/PI-negative cells, with single-stain controls used for compensation and gating. Four independent biological replicates (*n* = 4) were performed per cocktail and per differentiation stage. Among the four cocktails tested, VCCRI-ZyMix-4 consistently yielded the highest cell viability across stages and was therefore selected as the in-house Cardiomyocyte Dissociation Buffer for all downstream applications (see *hiPSC-Derived Cardiomyocyte In-house Dissociation Protocol*). Membrane and nuclear integrity following dissociation were further verified by cytospin preparation and confocal imaging of cTnT, VIMENTIN and DNA staining as described below.

### Functional and structural validation of hiPSC-derived cardiomyocytes by FluoVolt action potential recording and confocal imaging

To validate the structural and electrophysiological maturation of hiPSC-derived cardiomyocytes generated by our in-house directed differentiation protocol, day-12 hiPSC-CMs were dissociated using the *in-house Cardiomyocyte Dissociation Buffer* described below in *hiPSC-Derived Cardiomyocyte In-house Dissociation Protocol* and replated as monolayers on Matrigel-coated 12-well plate (Corning, USA) or on PhenoPlate 96-well, black, optically clear flat-bottom plates (Revvity) at a density of 100,000 cells/cm² in RPMI+B27+Glutamax supplemented with 200 µM L-Ascorbic acid and 5 µM Y-27632 (72304, STEMCELL Technologies). Replated cultures were maintained until day 22 post-differentiation, with media replaced every 2 days. For optical action potential recording, replated cardiomyocytes were loaded with 1× FluoVolt™ Membrane Potential Dye (F10488, Thermo Fisher Scientific) according to the manufacturer’s instructions and exchanged into phenol red-free RPMI 1640 (11835030, Thermo Fisher Scientific) supplemented to 1 mM CaCl2, as previously described [72]. In brief, cells were transferred to a Nikon Eclipse Ti2-E inverted microscope fitted with a Nikon Plan Fluor 10× objective (NA 0.3) and equilibrated in the live-cell imaging chamber at 37°C and 5% CO2 for 30 min prior to acquisition. Spontaneous action potentials were recorded using an Andor Zyla sCMOS high-speed camera (Oxford Instruments) at a 5 ms temporal resolution from 512 × 512-pixel regions of interest covering spontaneously contracting cardiomyocyte monolayers. Optical traces were obtained for 5-10 seconds and extracted in Excel. In parallel, structural maturation of replated cultures was assessed by fixation in 2% (v/v) PFA for 10 min, followed by immunofluorescence staining for cardiac troponin T (cTnT), α-actinin (ACTN2), and DNA, and confocal imaging on a Zeiss LSM900 as detailed in *Confocal Laser Scanning Microscopy* and *Immunofluorescence staining of cytospin preparations and replated cells* described in this Methods section.

### hiPSC-derived cardiomyocyte in-house dissociation protocol

Cells in 48-well plates were washed with 0.5 mL/well PBS and then incubated with 250 µL/well of room temperature *Cardiomyocyte Dissociation Buffer* for 30 minutes at 37°C, tapping the plate every 15 minutes [54]. The Cardiomyocyte Dissociation Buffer was composed of 12.5 U/mL Papain (Cat. #10108014001, Sigma Aldrich), 0.5 mM Cysteine, 132.5 U/mL Collagenase II (Cat. #LS004176, Worthington), 0.5% BSA (Cat. #A7906, Sigma), 0.025 mg/mL DNase I (Cat. #10104159001, Sigma Aldrich), and 0.25 U/mL Dispase (Cat. #07913, STEMCELL), all made up in DMEM (+L-Glutamine, +Sodium Pyruvate, +4.5 g/L Glucose). Following this, 500 µL FACS buffer/well was added, and the plate was placed on ice while still in the hood. Cells were gently pipetted up and down 20 times using a P1000 pipette to achieve single-cell status, with an additional 10 pipetting cycles if needed. The cells were then transferred to 15 mL Falcon tubes, pooling wells as required, and excess FACS buffer was added to each tube. The tubes were spun at 300 g for 4 minutes, and the supernatant was carefully removed. The cell pellets were resuspended by flicking. Cells were fixed in 1% PFA (prepared by diluting 4% PFA stock in FACS buffer) for 10 minutes. After fixation, excess PBS was added to the tubes, and they were spun down at 300 g for 4 minutes. The supernatant was removed, and the pellets were resuspended, ensuring no clumps were present. Finally, sufficient PBS (+0.02% Sodium Azide) was added to each tube, and the cells were stored at 4°C.

### Adult mouse cardiomyocyte isolation by gentleMACS™ Octo Dissociator (Miltenyi Heart Perfusion Kit)

For one of the two adult cardiomyocyte isolation strategies, hearts from 2- to 4-month-old wild-type C57BL/6J male and female mice were processed using the Miltenyi Heart Dissociation Kit, mouse and rat (130-098-373, Miltenyi Biotec) on the gentleMACS™ Octo Dissociator with Heaters (130-096-427, Miltenyi Biotec) according to the manufacturer’s instructions. Briefly, mice were sacrificed by cervical dislocation under deep anesthesia, hearts were rapidly excised, and the aorta was retrogradely perfused via the gentleMACS™ Heart Perfusion device with the kit perfusion buffer to flush residual blood and deliver the enzyme mix to the coronary vasculature. The perfused heart was then transferred into a gentleMACS™ C Tube (130-093-237, Miltenyi Biotec) containing pre-warmed Enzyme Mix and run on the gentleMACS™ Octo Dissociator using the manufacturer-supplied heart-dissociation program (37C_mr_Heart_01 followed by 37C_mr_Heart_02). The resulting cell suspension was filtered through a PluriStrainer (200 μm) into a 50 mL Falcon tube, washed in DMEM (+L-Glutamine, +Sodium Pyruvate, +4.5 g/L Glucose), and pelleted at 300 g for 5 min at room temperature. Pellets were resuspended in FACS buffer (PBS, 2% (v/v) FBS, 2 mM EDTA pH 7.9), counted, and processed for downstream applications including cytospin preparation, confocal microscopy and MNase digestion. Sample integrity was verified by confocal imaging of cytospun cardiomyocytes co-stained for α-actinin (ACTN2) and DNA, as detailed below.

### Adult mouse cardiomyocyte isolation by Langendorff retrograde perfusion

WT adult cardiomyocytes were rapidly isolated by Langendorff retrograde perfusion and purified by immunomagnetic cell separation as previously established by us [56]. Briefly, hearts from 3-month-old female C57BL/6J mice were cannulated via the aorta *in situ*, mounted on a Langendorff perfusion rig, and perfused with calcium-free perfusion buffer followed by an enzymatic digestion buffer. Atria were removed from the digested heart after perfusion and the ventricles were gently teased apart using forceps and resuspended into a single cell suspension by slow pipetting in transfer buffer and filtered. Cardiomyocytes were enriched by three low-speed centrifugations for 3 minutes at 20 x *g* to remove smaller non-myocytes. Further purification was performed using anti-CD31 antibody bound magnetic beads (details provided in [56]) to remove contaminating endothelial cells yielding a highly pure (∼95% cardiomyocytes) and viable (∼70%) cardiomyocyte preparation of ∼1.5 × 10□ cells per heart. Cardiomyocytes were fixed with 1% PFA (2 ml) for 10 minutes, washed by centrifugation at 100 x *g* for 4 minutes twice, resuspended in PBS (3 mL) and stored at 4°C. Cardiomyocyte purity and intact morphology were verified by flow cytometry using sequential gating for intact cells, DNA-positive (nucleated) events, and cTnT-positive (cardiomyocyte) events, as well as by confocal microscopy of cytospun preparations co-stained for ACTN2, cTnT and DNA. Cardiomyocytes prepared in this way (fixed in 1% PFA as described above) were used directly for MNase digestion (see *MNase digestion of adult mouse cardiomyocytes*), with cell or nucleus number adjusted as described below to account for the predominantly binucleated state of adult cardiomyocytes.

### Immunofluorescence staining of cytospin preparations and replated cells

Immunofluorescence staining of dissociated cells (cytospin preparations) and replated cardiomyocytes was performed using a saponin-based permeabilization workflow adapted from our previously published in-house protocol (Contreras et al., 2025). Briefly, 1% PFA-fixed cells deposited on Epredia™ SuperFrost Plus™ Adhesion slides by cytospin (see Cytospin of Cells in Suspension), or replated cardiomyocytes in Matrigel-coated glass-bottom dishes or PhenoPlate 96-well, black, optically clear flat-bottom plates (Revvity), were washed once with 1× PBS and permeabilized in 1× in-house permeabilization/wash buffer (0.2% (w/v) saponin, 4% (v/v) FBS or 2% (v/v) NCS, 1% (w/v) BSA, 0.02% (w/v) sodium azide in PBS) for 15 min at room temperature. After permeabilization, samples were incubated overnight at 4°C with primary or directly conjugated antibodies prepared in 1× permeabilization/wash buffer, in a humidified chamber to prevent evaporation. Antibodies used in this study included anti-cardiac troponin T (BD Pharmingen™ Alexa Fluor® 647 Mouse Anti-Cardiac Troponin T, BD Pharmingen™ PE Mouse Anti-Cardiac Troponin T; or unconjugated anti-cTnT, 1:200, MA5-12960, Thermo Fisher Scientific), Anti-α-Actinin (Sarcomeric) (clone REA402, Vio R667 conjugate, Miltenyi Biotec, REAfinity recombinant antibody), anti-VIMENTIN (1:100, D21H3, XP® Rabbit mAb, Alexa Fluor® 647 Conjugate, Cell Signaling Technology #9856), and species-appropriate Alexa Fluor-conjugated secondary antibodies (1:500, Thermo Fisher Scientific). The following day, samples were washed three times in 1× PBS, incubated with secondary antibodies (when required) in 1× permeabilization/wash buffer for 60 to 120 min at room temperature protected from light, and washed three times in 1× PBS. For mitochondrial network visualization in adult cardiomyocytes, MitoSpy™ Orange CMTMRos (424803, BioLegend) or MitoSpy™ Green FM (424805, BioLegend) was applied to fixed cardiomyocytes at 100 nM for 10 minutes. Nuclei were counterstained with a 1:5,000 dilution of Vybrant™ DyeCycle™ Violet (V35003, Thermo Fisher Scientific) in 1× permeabilization/wash buffer or with DAPI (D1306, Thermo Fisher Scientific) at 1 µg/mL in 1× PBS for 10 min at room temperature. Cytospin slides were mounted with Dako Fluorescence Mounting Medium (S3023, Agilent Technologies, Inc., USA) and #1.5 coverslips and stored at 4°C protected from light until imaging. Confocal imaging was performed on a Zeiss LSM900 as detailed before.

### MNase digestion of adult mouse cardiomyocytes

For chromatin profiling of adult mouse cardiomyocytes, isolated adult cardiomyocytes were aliquoted at 250,000 nuclei per tube, accounting for the ∼85% binucleation rate of adult cardiomyocytes (i.e. nucleus number rather than cell number was used for normalization) [56]. Tubes were made up to 1 mL with PBS and decrosslinked at 55°C for 10 min, then pelleted at 500 g for 5 min at 4°C. Pellets were resuspended in 200 μL Digestion Buffer #2 and processed as in the optimized workflow. For the time-course (Fig. 5F), five tubes were prepared (No-MNase, 1 min, 2.5 min, 5 min and 10 min). After CaCl2 supplementation, 2 μL of 1.5 U/μL MNase was added to the No-MNase, 1, 2.5 and 5 min tubes only, and tubes were placed on the Thermomixer at 37°C and 450 RPM with a 10 min master timer; the 5 min tube received MNase at the 5 min mark, the 2.5 min tube at the 7.5 min mark, and the 1 min tube at the 9 min mark, so that all reactions were stopped simultaneously at the 10 min endpoint with cold STOP Solution. For the concentration curve (Fig. 5G), six tubes were prepared (No-MNase, 0.5 U, 1 U, 3 U, 6 U and 10 U) using the MNase dilution series described above, and digested at 37°C and 450 RPM for 5 min. RNase A, Proteinase K, and GeneJET-based purification proceeded as described.

### Multi-parametric flow cytometry analyses

Multi-parametric flow cytometry analyses were performed using a BD LSRFortessaTM X-20 Cell Analyzer and a BD FACSymphony A5 High-Parameter Cell Analyzer, both equipped with five excitation lasers (UV 355 nm, Violet 405 nm, Blue 488 nm, Yellow/Green 561 nm, and Red 633 nm). FSC-H versus FSC-A, FSC-H versus FSC-W, and SSC-H versus SSC-W cytograms were used to discriminate and gate out doublets/cell aggregates during sorting or analysis. Data were collected using BD FACSDiva™ Software. For optimal DNA dye signal detection and cell cycle progression analyses, an event concentration of <1,000 events/seconds was used, and 25,000-50,000 events were captured for most cell types. For analysis of complex organs, including cells collected from embryonic and adult hearts, around 500,000-1,000,000 events were captured. All flow cytometry data were analyzed using FlowJo Portal (version 10.8.1, BD) using macOS Tahoe. Automated or manual compensation was done only when required.

### Confocal laser scanning microscopy

Confocal laser scanning microscopy was performed using a Zeiss LSM900 inverted microscope, which includes an upright Zeiss Axio Observer 7, Colibri 5 solid-state LED fluorescence light sources (solid-state laser lines: 405, 488, 561, 640), two Gallium Arsenide Phosphide photomultiplier tubes (GaAsP-PMT), and a motorized stage controlled by ZEN blue 3.4 software. Regular or tile images were acquired using objectives ranging from ×10 (0.45 NA with a WD 2.0, Air Plan-APO UV-VIS–NIR), ×20 (0.8 NA with a WD 0.55, Air Plan-APO UV-VIS–NIR), ×40 (1.3 NA with a WD 0.21, Oil Plan-APO DIC-UV-VIS–IR), and ×63 (1.2 NA with a WD 0.19, Oil Plan-APO DIC-UV-VIS–IR). The ×40 and ×63 objectives were used with immersion oil ImmersolTM 518 F (433802-9010-000, Zeiss). For z-stack 3D imaging, z-step sizes ranged from 0.25-1 μm, with images acquired under confocal settings using a motorized focus drive. Laser power during imaging was kept below 3.5%. When indicated, some tissue FFPE sections were imaged using a Leica Thunder Imager and Leica Image Files (LIFs) were imported directly in Fiji.

### Agarose gel electrophoresis of MNase-digested DNA

MNase-digested DNA was resolved on 2% to 2.5% (w/v) agarose gels prepared in 1× TAE buffer and stained with SYBR Safe DNA Gel Stain (S33102, Thermo Fisher Scientific). Gels were run at 90 V for 45 to 60 min alongside a 100 bp low-range DNA ladder (GWS-DMW-100L, GeneWorks) and imaged on a Bio-Rad ChemiDoc MP system. Mononucleosomal, dinucleosomal and trinucleosomal fractions were identified by their characteristic ∼150, ∼300 and ∼450 bp band positions [38]. Oligonucleosomes were defined as a smeared migration product lacking discrete banding, typically larger than 500 bp.

### Statistical analysis

For statistical analysis, unless otherwise specified, all results obtained from independent experiments are reported as means ± standard errors of means (SEM) of multiple replicates. Unless otherwise indicated, “*N*” in Figure Legends represents the number of animals or independent biological samples or replicates per group.

## AUTHOR CONTRIBUTIONS

C.T. Methodology, Validation, Formal Analysis, Investigation, Data Curation, Writing – Review and Editing.

D.T.H. Methodology, Validation, Formal Analysis, Investigation, Data Curation, Writing – Review and Editing, Visualisation.

M.N-S. Methodology, Validation, Formal Analysis, Investigation, Data Curation, Writing – Review and Editing, Visualisation.

A.M.N. Methodology, Validation, Writing – Review and Editing.

R.P.H. Validation, Writing – Review and Editing, Funding Acquisition.

O.C. Conceptualisation, Methodology, Validation, Formal Analysis, Data Curation, Writing – Original Draft Preparation, Writing – Review and Editing, Visualisation, Supervision, Project Administration, Funding Acquisition.

## Competing interests

The authors declare that they have no competing interests.

**Correspondence** and requests for materials and methods should be addressed to Osvaldo Contreras.

## Acknowledgments

We are grateful to Bernice Stewart for administrative support and to the Victor Chang Cardiac Research Institute Innovation Centre (funded by the New South Wales Government Ministry of Health) for infrastructure support. We also thank the Garvan Weizmann Center for Cellular Genomics (GWCCG) and the Australian Genome Research Facility (AGRF) for library QC and next-generation sequencing, and the Core Facility Flow at the Garvan Institute of Medical Research (GIMR) for flow cytometry support. O.C. acknowledges support from the Medical Advances Without Animals Trust (MAWA), an organisation dedicated to advancing medical science and improving human health and therapeutic interventions without the use of animals or animal products. We thank Jordan Thorpe for initial guidance with FluoVolt™ membrane potential recordings, Profs. Robert (Bob) Graham, Adam Hill, and Sally Dunwoodie for their mentorship and generosity, and Marion Lingard and Matt Banfield from Miltenyi Biotec for facilitating access to the gentleMACS™ Octo Dissociator with Heaters. Figures were created using Adobe Illustrator and Adobe Photoshop 2026. We also thank Felipe Serrano (www.illustrative-science.com) for help with the design of Figure 6.

## Funding

This work was supported by the National Health and Medical Research Council (NHMRC) of Australia (Investigator Grant L3 GNT2008743 and Senior Principal Research Fellowship GNT1118576 to R.P.H.) and the Australian Research Council (DP210102134 to R.P.H.). M.N-S. was supported by the National Health and Medical Research Council (NHMRC) of Australia (Investigator Grant EL1 Grantor Reference ID 2033823). O.C. was supported by the Medical Research Future Fund (MRFF) Stem Cell Therapies Mission (2024/MRF2032746); the Miltenyi Research Award 2022; a joint Medical Advances Without Animals Trust (MAWA) and Rozzoli Family Fund (RFF) Research Grant; and a 2024 NSW Cardiovascular Research Network (CVRN) and Victor Chang Cardiac Research Institute (VCCRI) Innovation Grant, co-funded by the National Heart Foundation of Australia and the Office for Health and Medical Research (OHMR), New South Wales Government. This work was also supported by the NSW Health Cardiovascular Research Capacity Program (2026 to 2029) through a Cardiovascular Early- to Mid-Career Researcher Grant to O.C.

## Data and materials availability

All data needed to evaluate the conclusions of the paper are present in the main manuscript and/or the Supplementary Materials. Raw datasets will will be deposited prior to publication at the NCBI Gene Expression Omnibus (GEO) database.

## Notes

### Competing Interest Statement

The authors have declared no competing interest.

