## Supplemental Figure 1 for "A scalable MNase-seq framework for reproducible nucleosome profiling across pluripotent stem cell and cardiomyocyte models"

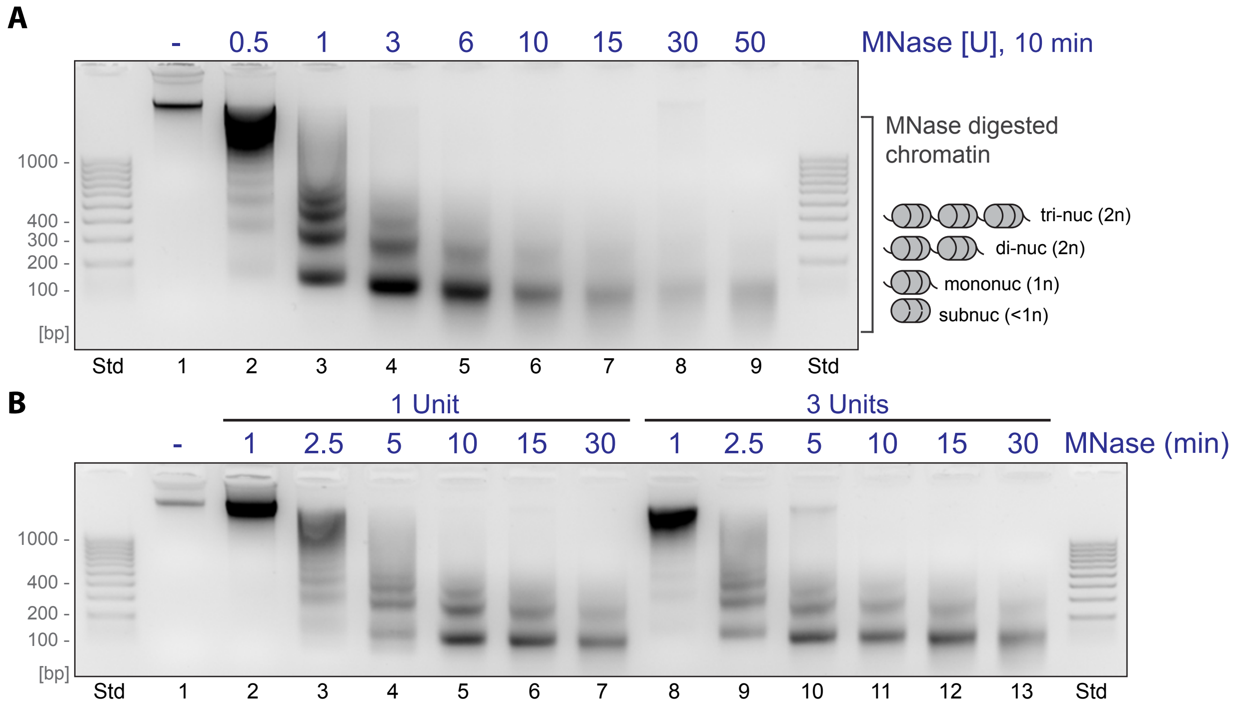


**Supplementary Figure 1. MNase titration and time-dependent MNase-mediated digestion of chromatin in hiPSCs.** (A) 2.5% agarose gel electrophoresis of MNase digestion products from a typical MNase titration experiment. The concentration-dependent MNase-digested chromatin gel displays the characteristic "nucleosome ladder" pattern, with bands corresponding to mono-nucleosomes (1n), di-nucleosomes (2n), tri-nucleosomes (3n), and longer oligonucleosome fragments. (B) 2.5% agarose gel electrophoresis of MNase digestion products comparing two MNase units (1 vs. 3) in a time-dependent experiment. The time-dependent MNase-digested chromatin exhibits the characteristic "nucleosome ladder" pattern described above. 500,000 hiPSCs were used in A and B.
