## Supplemental Figure 2 for "A scalable MNase-seq framework for reproducible nucleosome profiling across pluripotent stem cell and cardiomyocyte models"

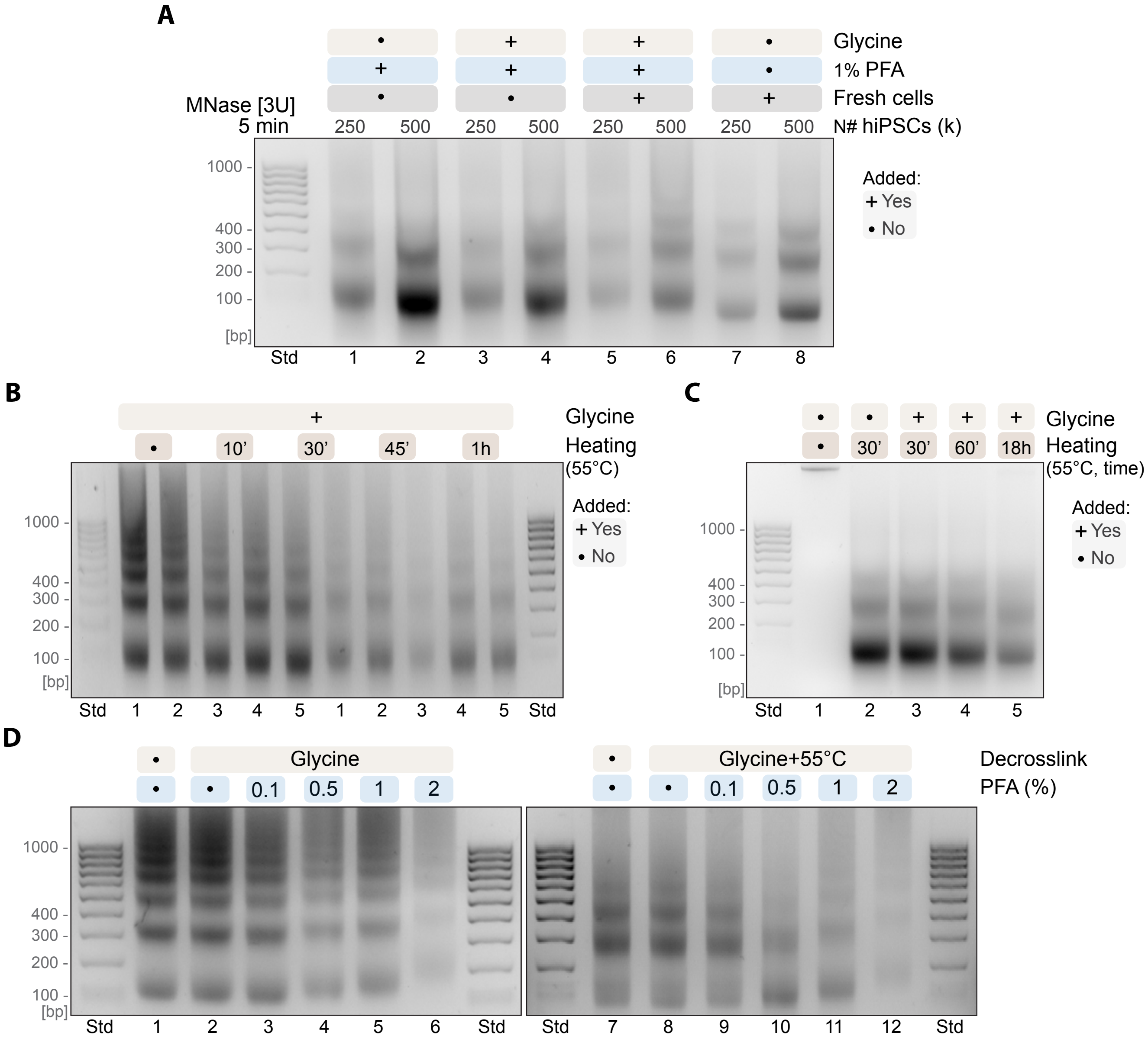


**Supplementary Figure 2. Optimization of PFA fixation and crosslink reversal for MNase-mediated chromatin digestion.** All panels show MNase-digested hiPSC chromatin resolved by agarose gel electrophoresis (2.5% w/v agarose). Thermal decrosslinking refers to heating fixed cells at 55°C to reverse PFA crosslinks prior to digestion; equal volumes of purified DNA were loaded in each lane. (A) Storage stability of fixed chromatin. Nucleosomal ladders from 1% PFA-fixed hiPSCs (10 min fixation) stored for 2–4 weeks at 4°C are compared with freshly dissociated hiPSCs processed in parallel, both fixed and unfixed. Samples were digested with 3 U MNase (5 min, 37°C) at inputs of 250,000 and 500,000 cells. (B–C) Effect of thermal decrosslinking on fixed chromatin at two digestion times. 1% PFA-fixed hiPSCs (10 min fixation) stored for 1 week were heated at 55°C for increasing durations to reverse crosslinks before MNase digestion (3 U), and nucleosomal resolution was compared across heating conditions. 20 µL of purified DNA loaded per lane; 400,000 cells per condition, except lane 1 (No-MNase genomic DNA control, 175,000 cells). (B) 2.5 min digestion. (C) 5 min digestion. (D) Synergy of chemical quenching and thermal decrosslinking across PFA concentrations. Freshly dissociated hiPSCs (250,000 cells per condition) were fixed on ice for 10 min with 0.1%, 0.5%, 1%, or 2% PFA (plus no-PFA and no-PFA/no-quench controls), then either quenched with glycine alone or quenched and heated at 55°C for 30 min, before digestion with 3 U MNase (2.5 min, 37°C). 20 µL of purified DNA loaded per lane.
