## Supplemental Figure 3 for "A scalable MNase-seq framework for reproducible nucleosome profiling across pluripotent stem cell and cardiomyocyte models"

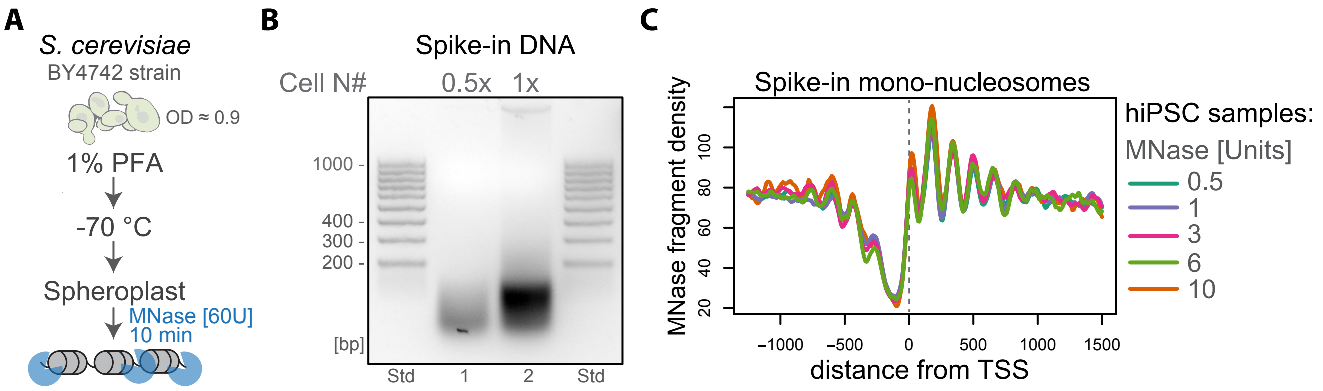


**Supplementary Figure 3. Custom-generated spike-in DNA from log-phase growing yeast.** (A) Schematic illustrating the in-house development of spike-in DNA from growing yeast. OD, optical density. (B) Spike-in DNA was generated using 60U of MNase for 10 minutes with two different cell quantities. The 1x cell quantity was selected for downstream spike-in analysis as it contained higher and more abundant molecular weight DNA. (C) Yeast MNase-seq DNA fragment density relative to the distance from the transcriptional start site (TSS) using increasing concentrations of MNase (0.5U to 10U), demonstrating similar footprinting patterns of spike-in mono-nucleosomes.
