## Supplemental Figure 4 for "A scalable MNase-seq framework for reproducible nucleosome profiling across pluripotent stem cell and cardiomyocyte models"

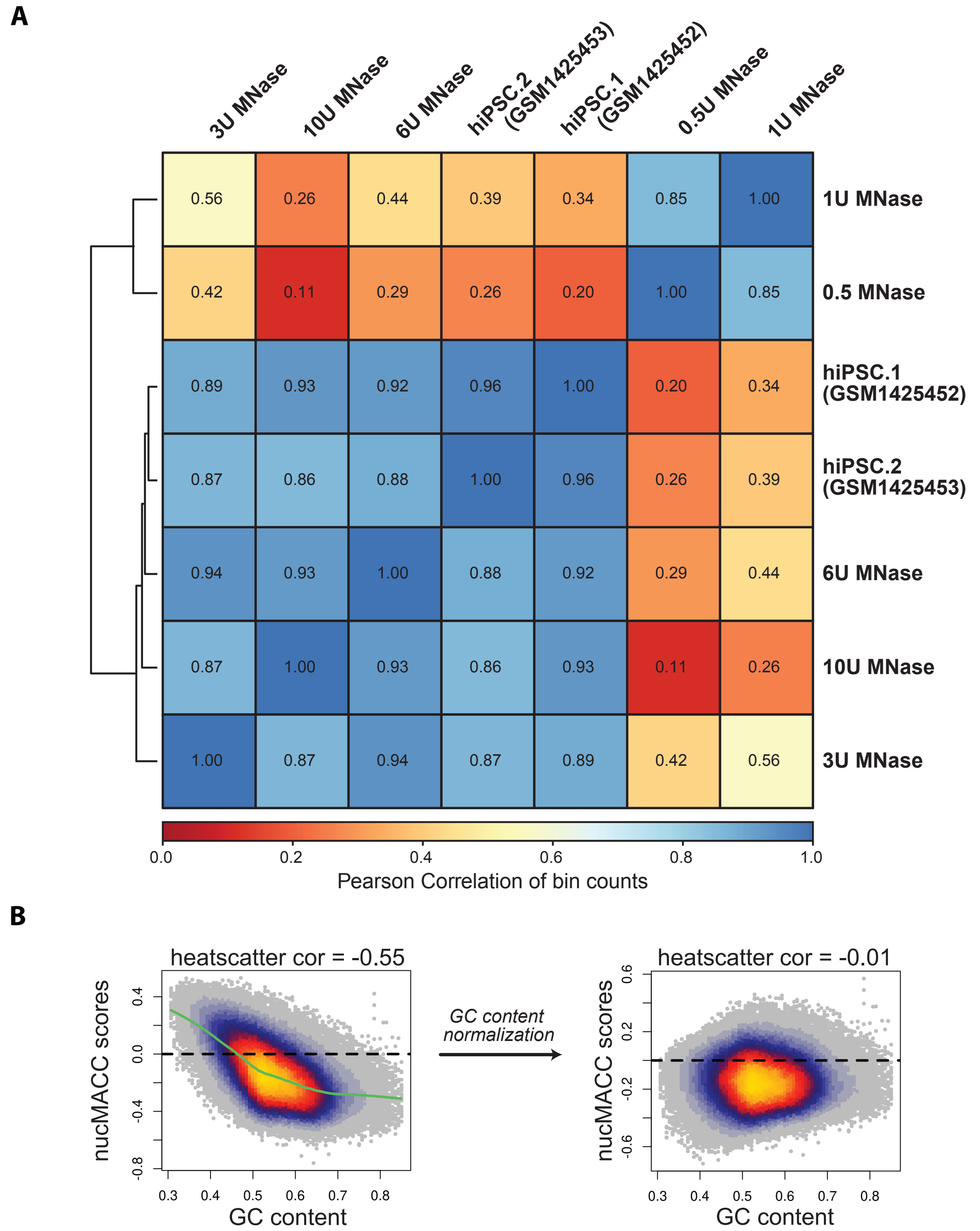


**Supplementary Figure 4. Pearson correlation of mononucleosomal read counts across MNase titration conditions and published MNase-seq datasets.** (A) Heatmap showing pairwise Pearson correlation coefficients of genome-wide mononucleosomal read counts (bin counts) across five MNase titration conditions (0.5, 1, 3, 6, and 10 U) generated in this study and two replicates from a published hiPSC MNase-seq dataset [West et al., 2014; NIH GEO database under accession code GSE59064]. Hierarchical clustering was performed on correlation values. Higher MNase concentrations (3–10 U) cluster with the published datasets (r = 0.86–0.93), indicating high concordance between optimized digestion conditions and independent reference data. Lower MNase concentrations (0.5 and 1 U) form a separate cluster with markedly reduced correlation coefficients relative to both higher MNase conditions and published data (r = 0.20–0.39), consistent with incomplete chromatin digestion at these concentrations. Colour scale represents Pearson correlation from 0.0 (red) to 1.0 (blue). (B) Raw nucMACC scores show a strong negative correlation with nucleosome GC content (heatscatter correlation = –0.55), reflecting the concentration-dependent AT cleavage preference of MNase. Following LOESS-based GC content normalisation, this bias is effectively removed (heatscatter correlation = –0.01), ensuring that nucMACC scores reflect chromatin accessibility independently of local sequence composition.
