## Supplemental Figure 5 for "A scalable MNase-seq framework for reproducible nucleosome profiling across pluripotent stem cell and cardiomyocyte models"

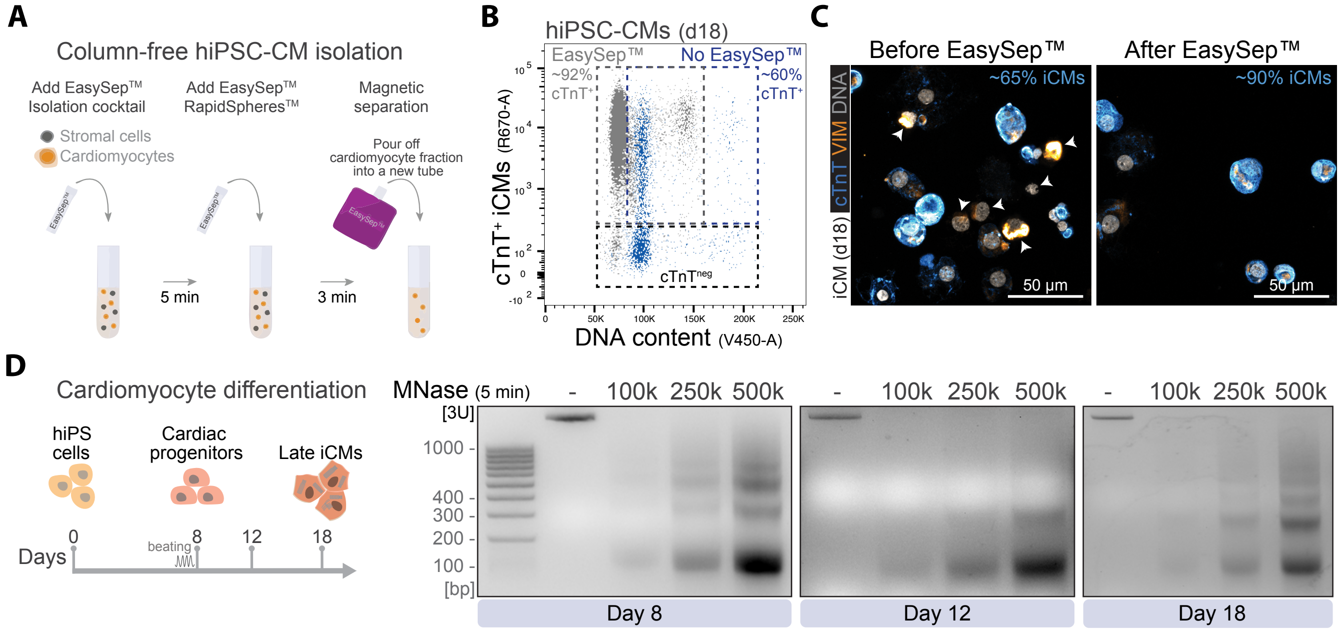


**Supplementary Figure 5. Characterization and magnetic cardiomyocyte enrichment using EasySep™.** (A) Schematic illustrating the EasySep™ column-free hiPSC-derived cardiomyocyte isolation. The EasySep™ Human PSC-Derived Cardiomyocyte Enrichment Kit isolates hiPSC-derived cardiomyocytes from cell cultures by negative selection. Unwanted cells are targeted for removal with tetrameric antibody complexes recognizing non-cardiomyocytes and dextran-coated magnetic particles. (B) Flow cytometry plot showing the EasySep™ hiPSC-CM-enriched fraction, totaling ~92% of cTnT+ cells vs no EasySept™ separation (n = 3). (C) Confocal microscopy images of cytospun hiPSC-CM (iCM) day 18 cultures, with cTnT, VIMENTIN (VIM) and DNA staining (n = 2). Arrows indicate non-cardiomyocyte stromal cells. (D) MNase digestion products were analyzed using different hiPSC-CM cell numbers at day 8, 12, and 18 of directed differentiation. Undigested genomic DNA (lanes noted as "-") was purified from 100,000 (100k) hiPSC-cardiomyocytes .
